# Rif1 regulates telomere length through conserved HEAT repeats

**DOI:** 10.1101/2020.11.24.394866

**Authors:** Calla B Shubin, Rini Mayangsari, Ariel D Swett, Carol W Greider

## Abstract

In budding yeast, Rif1 negatively regulates telomere length, but the mechanism of this regulation has remained elusive. Previous work identified several functional domains of Rif1, but none of these has been shown to mediate telomere length. To define Rif1 domains responsible for telomere regulation, we localized truncations of Rif1 to a single specific telomere and measured telomere length of that telomere compared to bulk telomeres. We found that a domain in the N-terminus containing HEAT repeats, Rif1_177-996_, was sufficient for length regulation when tethered to the telomere. Charged residues in this region were previously proposed to mediate DNA binding. We found that mutation of these residues disrupted telomere length regulation even when Rif1 was tethered to the telomere. Mutation of other conserved residues in this region, which were not predicted to interact with DNA, also disrupted telomere length maintenance, while mutation of conserved residues distal to this region did not. Our data suggests that conserved amino acids in the region from 436 to 577 play a functional role in telomere length regulation, which is separate from their proposed DNA-binding function. We propose that the Rif1 HEAT repeats region represents a protein-protein binding interface that mediates telomere length regulation.

## INTRODUCTION

Telomeres contain repetitive DNA that protects the ends of linear chromosomes in eukaryotes and allows for telomere length maintenance. Telomeres shorten due to the end replication problem. To counteract this shortening, telomerase adds telomere repeats and establishes an equilibrium length, which varies by species. Telomere binding proteins help maintain this equilibrium by positively or negatively regulating telomere addition (1,2). When the length equilibrium is disrupted, short telomeres can signal a DNA damage response, resulting in cellular senescence or cell death (3). Maintenance of telomere length equilibrium is critical for human health; short telomeres can cause age-related degenerative diseases, including pulmonary fibrosis, immune deficiency, and bone marrow failure; conversely, long telomeres lead to a predisposition to cancer (4,5). Identifying the mechanistic basis of telomere length regulation is therefore important to understand the role of telomeres in disease. Here we focus on the protein Rif1, which regulates telomere length equilibrium in yeast.

*Saccharomyces cerevisiae* Rif1 is a 1,916 amino acid protein, which regulates several processes, including origin firing, DNA repair, and telomere length. Rif1 was discovered in yeast through its interaction with the telomere binding protein Rap1. Deletion of *RIF1* results in long telomeres, indicating that it negatively regulates telomere length (6). Rif1 has domains that bind PP1 (Protein Phosphatase 1), Dbf4 (the regulatory component of DDK), and Rap1 (Figure 1A). Yeast Rif1 binds to PP1 via two canonical PP1 binding motifs, RVxF and SILK, in its N-terminus. This function is conserved from yeast to mammals, but, in mammalian Rif1, these binding sites are located in the C-terminus. The Rif1-PP1 complex blocks origin firing by de-phosphorylating Mcm4 in the pre-replication complex (7–11). We recently showed that this conserved Rif1 function does not regulate telomere length in yeast (12). The Rif1 C-terminus binds to Dbf4 (7,8), and also localizes Rif1 to the telomere through the Rap1 binding motif (RBM) (13), but deletion of these domains does not lead to telomeres that are as long as *rif1Δ* (12,13). Thus, while we know several binding partners of Rif1, the domains and mechanism by which Rif1 negatively regulates telomere length in yeast remain elusive.

**Figure 1:**
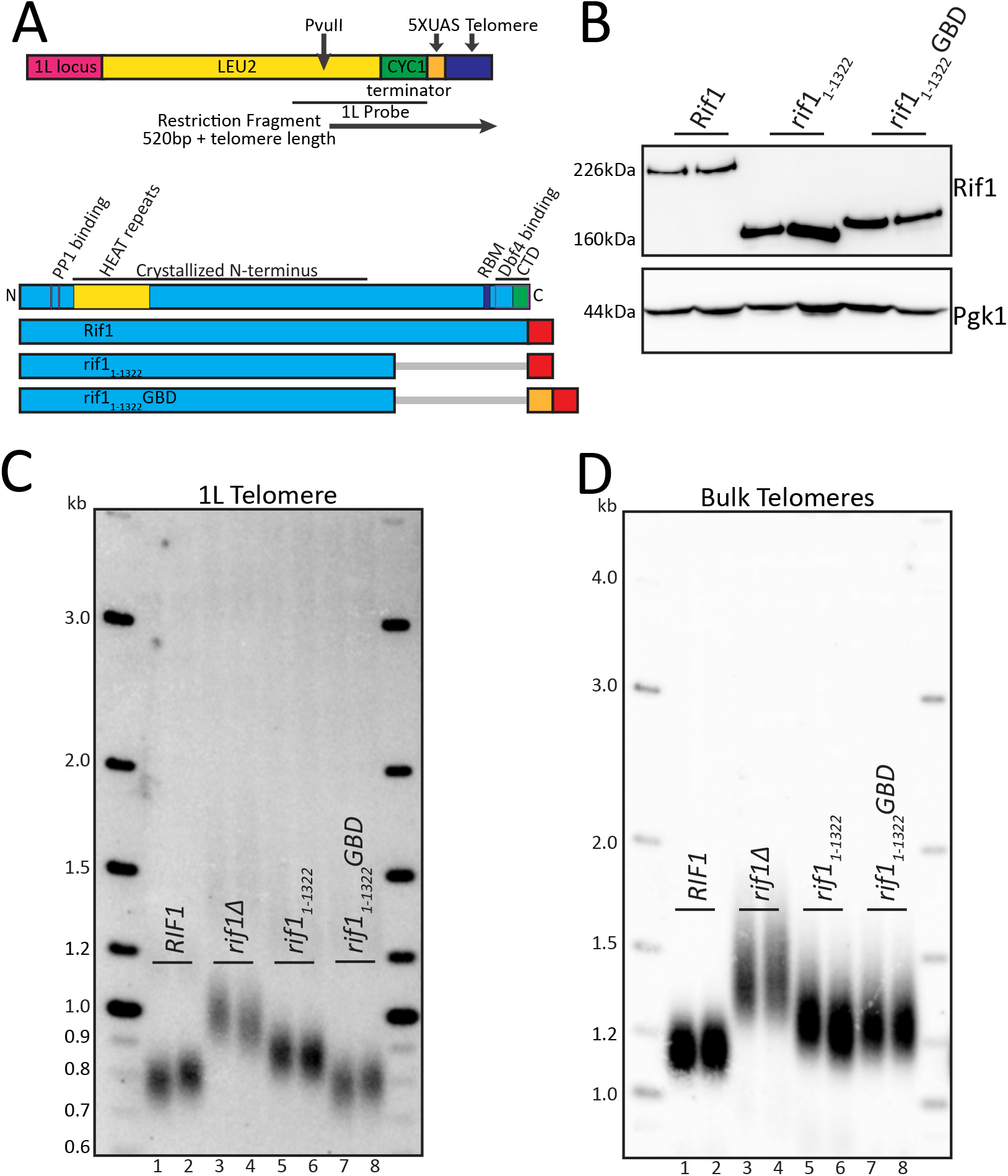
Rif1 N-terminus when localized to telomere is functionally sufficient in telomere regulation. A. Diagram of 5XUAS landing pad and evaluated Rif1 constructs. PvuII digestion of genomic DNA was used to detect the 5XUAS telomere length by Southern blot with a unique 1L telomere probe, the sequence of which is indicated by the black bar beneath the diagram. The 1L telomere restriction fragment is indicated by the black arrow below the diagram beginning at the PvuII cut-site. The lower section of the diagram shows a Rif1 domain map depicted to scale (RBM: Rap1 binding motif; CTD: Carboxyl-terminal domain; Dbf4 binding overlaps CTD). Below is a schematic of the Rif1 constructs tested in this figure (Blue: Rif1; Red: 6xFLAG; Orange: GBD; Grey bar: C-terminal truncation of residues). B. Western blot showing Rif1 (anti-FLAG antibody) and Pgk1 (anti-Pgk1 antibody, control) protein levels of indicated strains. C. Southern blot showing 1L telomere probe for the indicated strains. D. Southern blot from C, rehybridized to show bulk Y’ telomere probe for the indicated strains.

Rif1 also plays a role in the DNA damage response in regulating non-homologous end joining (NHEJ). Rif1 residues implicated in DNA binding are important for carrying out Rif1 NHEJ function (14). These residues are located in the most conserved region of Rif1 homologues (15). In mammalian cells, Rif1 is recruited to double-stranded breaks (DSB) by phosphorylated 53BP1 (Rad9 in yeast). ATM (Tel1 in yeast) phosphorylation of 53BP1 is required for recruitment of Rif1, which subsequently suppresses 5’ end resection at double-strand breaks (DSB). This function counteracts BRCA1 mediated homologous recombination at DSB (16–19). The precise mechanisms by which Rif1 promotes NHEJ in yeast, and whether these functions are related to those in telomere length regulation, are not yet understood.

Since the domains of Rif1 defined to date do not explain its role in regulating telomere length, we set out to identify a telomere functional domain. We tethered several independent domains of Rif1 to a unique telomere and measured telomere length at that telomere and also across the bulk telomere population. We found that one conserved region of HEAT repeats in the Rif1 N-terminus, aa 177-996, is sufficient to maintain telomere length regulation when tethered to the unique telomere. Mutational analysis suggests that residues within this domain, aa 436-577, may represent a protein-binding interface that promotes telomere regulation.

## MATERIALS AND METHODS

### Reagents

Enzymes used for cloning and Southern blots are all listed in Supplementary Table 1. Antibodies used are indicated in Western blotting below.

### Biological Resources

Yeast strains and plasmid vectors are all listed in Supplementary Table 1 and available upon request. Bacteria used for cloning are also listed in Supplementary Table 1.

### Computational Resources

N/A

### Statistical Analyses

N/A

### Molecular cloning

As described in (12). Plasmid and strain design were done *in silico* using SnapGene software. Standard molecular biology techniques including PCR and Gibson assembly (New England Biolabs) were used to make all plasmids and homology repair constructs for all yeast transformations. All constructs used to generate strains, including plasmids and oligos, as well as enzymes used for digest before transformation, are listed in Supplementary Table 1. All plasmid maps are available upon request. All restriction enzymes used in these studies are from New England Biolabs. All oligonucleotides and gene blocks were ordered from Integrated DNA Technologies (idtdna.com).

### Gal4 DNA binding domain (GBD) 5XUAS 1L telomere

The Gal4 DNA binding site construct was designed *in silico* using SnapGene software and constructed using standard cloning techniques. Homology arm for one-ended recombination was chosen using BlastN (https://blast.ncbi.nlm.nih.gov/) to determine a unique site on chromosome arm 1L upstream of the X-element and other duplicated genes (CS218, Supplementary Table 1). *LEU2* was used as a selectable marker, with a silent change T1674A to create a novel PvuII cut-site for telomere Southern analysis. *CYC1* terminator was included to prevent *LEU2* transcription into the 5XUAS landing pad. The 5XUAS binding site was inserted directly next to a 40bp telomere seed sequence with an I-Sce1 site, which, once cut, telomerase can recognize and extend, leading to *de novo* telomere elongation *in vivo* (20). PCR was used to amplify the construct from pCS33 while simultaneously adding 40bp of homology (CS155 and CS218, Supplementary Table 1) to chromosome 1L for one-ended homologous recombination, and then I-Sce1 digest was used to create the 3’ overhang before standard yeast transformation as described below. Transformed yeast were plated on minimal media plates lacking leucine, supplemented with nicotinamide (NAM) at 5mM. NAM allows for selection of inserted genes in heterochromatic regions of the genome such as the telomere by counteracting Sir2 chromatin modifications to allow expression of the selectable marker (21).

### Yeast culturing and transformation

Cells were grown logarithmically in yeast peptone dextrose (YPD) to an optical density of close to 1.0 described in (12). Cells were washed and resuspended in sterile water and 0.1M Lithium acetate (LiAc, Sigma L6883-250G) before pelleting. 50μL of the cell pellet was used in the transformation alongside the DNA homology repair template, 0.1M LiAc. Herring sperm DNA was added as a carrier for most transformations. The transformation reaction was incubated at 30°C for 10 minutes before adding 500μL Polyethylene Glycol (PEG, Sigma P4338-1KG) and another incubation at 30°C for 30 minutes. The heat shock step was performed at 42°C for 15 minutes to 1 hour. Cells were washed by adding sterile water to bring the volume up to 1mL before pelleting; the wash step was often repeated with another 1mL sterile water before plating. If selecting for a drug resistant marker such as KANMX, cells were resuspended after the wash step in 1mL YPD and recovered at 30°C for 4 hours. Oligonucleotides or enzymes used to amplify and isolate the homology repair template are listed with the strain list in Supplementary Table 1.

### Southern blotting

Genomic DNA (gDNA) was isolated by pelleting 1.5mL saturated overnight yeast culture grown in YPD, then beating the cells using 0.5mm glass beads in 250μL lysis buffer and 200μL phenol chloroform. Cells were spun down for 10 minutes at 14,000rpm, and ~200μL of the top clear solution, containing the gDNA, was carefully taken out. gDNA was precipitated using 500μL of 95% ethanol and then pelleted. The supernatant was discarded, and 500μL of 70% ethanol was added. Pellets were left to dry before resuspension in 50μL TE with RNaseA for 1 hour at 37°C or overnight at 4°C. 10μL of gDNA was digested using PvuII and XhoI for 1-4hours at 37°C. Of note, PvuII and XhoI were both used together to digest genomic DNA for all Southern blots to visualize the 1L telomere (PvuII) and bulk telomere (XhoI) restriction fragments. To better visualize the 1L telomere, 10ng of 2-log ladder (NEB N3200L) was loaded instead of 100ng, as done previously (12). Digested gDNA and ladder were loaded onto a 1% agarose gel and electrophoresed overnight at ~47V in 1XTTE. The gel was denatured for 30 minutes in a rocking shaker (1.5M NaCl, 0.5M NaOH) and was neutralized for 15 minutes (1.5M NaCl, 0.5M Tris, pH 7.0). gDNA on the gel was vacuum transferred onto a hybond nylon membrane (GE Healthcare GERPN303B) with 10XSSC (1.5M NaCl 0.17M NaCitrate, dihydrate) and UV cross-linked before blocking in Church buffer (0.5M Na2HP04, pH7.2, 7% SDS, 1mM EDTA, 1% BSA) for ~1 hour at 65°C. ^32^P radiolabeled PCR fragments were added onto the membrane and left to incubate overnight. 1L telomere Southern blots were hybridized with a radiolabeled purified PCR product of the *LEU2* gene and *CYC1* terminator amplified from a plasmid (pCS206) using primers CS414 and CS432, and bulk telomere Southern blots were hybridized with a Y’ PCR product. Oligonucleotide sequences used to generate the PCR products are listed in Supplementary Table 1. The membrane was washed 3-4 times for 15 minutes each, with 1XSSC 0.1%SDS buffer before laying down a phosphor screen (GE Healthcare). 1L telomere Southern blots were exposed to phosphor screens for 4-5 days and Y’ telomere Southern blots were exposed for 1 day. Images were captured on a STORM using ImageQuant software (GE Healthcare), and the .gel files were copied into PowerPoint and saved as .tif files. Telomere length was measured from at least two clones of each genotype.

### Western blotting

Protein extraction using trichloroacetic acid (TCA, Sigma T0699) and western blotting methods are similar to those described in (12). 500μL of cells were collected from overnight saturated yeast cultures grown in YPD. Cells were pelleted, and 1:10 ratio of TCA to water was added. Tubes were inverted to gently mix and left to incubate at room temperature for 30 minutes. Cells were pelleted and resuspended in 500μL 1M HEPES (Teknova H1035), and then centrifuged to remove the supernatant. Pellets were resuspended in 50μL 2X LDS sample buffer (Invitrogen NP0007), supplemented with 100mM DTT, and vortexed with ~50μl of 0.5mm glass beads. 50μl of LDS buffer was added, and the sample was boiled at 100°C for 5 minutes, and then spun down for 10 minutes, before carefully taking the supernatant. Samples were kept on ice or −20°C freezer until ready to be used, and re-boiled right before loading onto the gel. 3-8% Tris-Acetate gels (Invitrogen EA0375) were used to resolve all proteins, run at 150V for 1 hour and 20 minutes. Short transfer using Trans-Blot Turbo transfer system (Bio-Rad) with the pre-set 10-minute-high MW program was used for all westerns unless blotting for full length Rif1, where a long transfer using NuPAGE XCell II Blot Module (Thermofisher/Invitrogen EI9051) was used at 30V for 1.5 hours. Both αFLAG for Rif1 blots (1:1000) (Sigma M8823) and αPgk1 (1:10,000) (Invitrogen 459250) antibodies were blocked in 5% milk TBS-T (1X TBS 0.1% Tween-20), and secondary for both was αMouse (1:10,000) (Bio-Rad 1706516) also in 5% milk TBS-T. Forte HRP substrate (Millipore WBLUF0100) was used for imaging FLAG blots, and SuperSignal West Pico PLUS Chemiluminescent Substrate (Thermo 34580) was used for imaging Pgk1 blots. Signals were captured on an LAS-4000 imager (GE healthcare). Images were visualized using ImageQuant software (GE Healthcare), and the .gel files were copied into PowerPoint and saved as .tif files. Rif1 protein levels were visualized by western blot in at least two clones of each genotype.

### Protein Stability

Several constructs, all expressed at the endogenous *RIF1* promoter, had unexpected changes in Rif1 protein levels that limited data interpretation. We tested a construct containing only the C-terminal region, from amino acids 1323-1916, *rif1-NLS_1323-1916_*GBD. Unexpectedly, this construct was highly overexpressed even at the endogenous *RIF1* promoter, and ran at a much higher than expected molecular weight. We found a partial rescue of telomere length at both the 1L telomere and at bulk telomeres in this strain. Disrupting the Rap1 binding motif with a two amino acid substitution, I1762R and I1764R (13), still showed partial rescue, indicating binding to Rap1 may not be required for this C-terminal construct. While it is difficult to make a conclusion due to the high level of expression, we cannot rule out some role of the C-terminal region in regulating telomere length. In addition, we generated eight constructs containing internal deletions in Rif1 (Δ1-152, Δ191-340, Δ342-484, Δ486-684, Δ686-893, Δ895-1142, Δ1144-1221, Δ1223-1318). We found most of these constructs had low to undetectable protein levels on western blots, precluding this approach for domain mapping.

### Nuclear localization signal (NLS) identification

Potential NLS were identified using nls-mapper.iab.keio.ac.jp, using *S. cerevisiae* RIF1 protein sequence and using a strong cut-off parameter of 7 to identify NLS consensus. The only strong predicted NLS was located at position 56 with a high score of 13.

### Structure guided sequence alignment

Structure-guided sequence alignment was generated by first employing a multiple sequence alignment (MSA) of RIF1 orthologs from 12 yeast species using Clustal Omega (http://www.clustal.org/omega/). A FASTA file that contained a compilation of the RIF1 protein sequence, obtained from Orthogroup Repository Documentation and UniProt database, was used in generating the multiple sequence alignment. The output Clustal Omega file was then uploaded to the ConSurf Server (https://consurf.tau.ac.il) with the crystal structure of the RIF1-NTD dimer in complex with DNA double helicase (PDB: 5NW5) as a reference and S. cerevisiae as the query sequence. The ConSurf Server generated a PDB file in which the residues were color coded based on their conservation score ranging from 1-9, with the score of one being the most variable and nine being the most conserved. The RIF1 structure-guided alignment PDB file was then visualized and analyzed in PyMOL (The PyMOL Molecular Graphics System, Version 2.2.0, Schrödinger, LLC) (Supplementary Figure S1D). We also visualized the alignment using SnapGene software (Supplementary File 1) and graphed the conservation scores using GraphPad Prism 5.

## RESULTS

### The C-terminus of Rif1 is not required for telomere length regulation

To examine the regions of Rif1 that mediate telomere length regulation, we generated a yeast strain in which we could localize domains of Rif1 directly upstream of a unique telomere. We introduced 5 copies of the GAL4 <u>u</u>pstream <u>a</u>ctivating <u>s</u>equence, 5XUAS, immediately adjacent to telomere repeats on the left arm of chromosome 1 (1L) by eliminating the 1L subtelomeric X element (Figure 1A; Materials and Methods). We then fused the <u>G</u>AL4 DNA <u>b</u>inding <u>d</u>omain, GBD, to domains of Rif1 to localize them to this unique telomere (Figure 1A,B; Materials and Methods). We analyzed the 5XUAS 1L telomere by Southern blot with a unique probe to this region (see Materials and Methods). The 1L telomere showed a length distribution with a midpoint of 800bp, which includes the telomere and ~500bp of subtelomere, indicating its length distribution is regulated similarly to wild type yeast telomeres (Figure 1C, lanes 1-2). Deletion of *RIF1* resulted in elongation of the 1L telomere and bulk telomeres, as expected (Figure 1C, lanes 3-4; Figure 1D, lanes 3-4).

To begin dissecting Rif1 domains, we initially deleted the Rap1 binding site using a C-terminal truncation, *rif1_1-1322_*, which removes both Rap1 binding and Dbf4 binding (Figure 1A,B). This construct showed long telomeres at 1L, indicating loss of telomere length regulation (Figure 1C). When the same Southern was re-hybridized with the subtelomeric Y’ probe to visualize most other telomeres (“bulk telomeres”, see Materials and Methods), *rif1_1-1322_* bulk telomeres were also long. Notably, both the 1L and bulk telomeres were not as long as *rif1Δ* (Figure 1C compare lanes 5-6 to 3-4; Figure 1D compare lanes compare lanes 5-6 to 3-4). This partial effect of *rif1_1-1322_* may be due to a lower concentration of Rif1 near the telomere when it is not bound to Rap1.

To control recruitment of Rif1 to the 1L telomere, we fused the GBD to *rif1_1-1322_*, to create *rif1_1-1322_GBD* at the *RIF1* genomic locus. Strikingly, *rif1_1-1322_GBD* restored telomere length at 1L to essentially wild type length (Figure 1C compare lanes 7-8 to 1-2). This result shows that Rif1 residues 1-1322 are fully functional when recruited to a single telomere. This rescue effect was not seen at bulk telomeres, which were long, as expected, since *rif1_1-1322_*GBD is not recruited to bulk telomeres (Figure 1D compare lanes 7-8 to 5-6 to 3-4). This data indicates that the N-terminal region of Rif1, when tethered to the telomere, is sufficient for telomere length regulation. We further used this platform to probe Rif1 functional domains by comparing the effects on the single 1L telomere, to which Rif1-GBD can bind, to the global effects on bulk telomeres.

### Identification of a required nuclear localization signal in the N-terminus of Rif1

Having defined the N-terminus of Rif1 as sufficient for telomere regulation when tethered by the GBD, we next sought to narrow down this important region. We previously demonstrated that point mutations in the N-terminal PP1 binding site (aa 114 to 149) disrupted origin firing but were dispensable for telomere length regulation (12). To further probe this region, we removed 176 amino acids from the N-terminus, which we refer to as the N-terminal domain, or NTD. This NTD deletion removes the PP1 binding sites (Figure 2A). The Rif1_177-1322_GBD construct (Figure 2A) was stably expressed, and, in fact, had somewhat higher steady state levels than rif1_1-1322_GBD, as indicated by western blot (Figure 2B). We found that 1L telomeres were elongated in *rif1_177-1322_GBD* cells, in spite of this increased protein expression (Figure 2C, lanes 9-10), indicating that the NTD has a functional role in telomere regulation. To examine this result further, we used computational analysis and found a predicted Nuclear Localization Signal (NLS) in this region, beginning at amino acid 56 (see Materials and Methods). To promote nuclear localization of the fusion protein, we added c-myc NLS (22) to generate *Rif1-NLS_177-1322_GBD* (Figure 2A,B). This construct restored the telomere length distribution of telomere 1L to that of wild type (Figure 2C compare lanes 11-12 to 1-2). However, the bulk telomeres were still long, as expected, since the construct was not localized to bulk telomeres (Figure 2D, lanes 11-12 and 9-10). This data suggests that the NTD delivers a critical NLS for Rif1 function.

**Figure 2:**
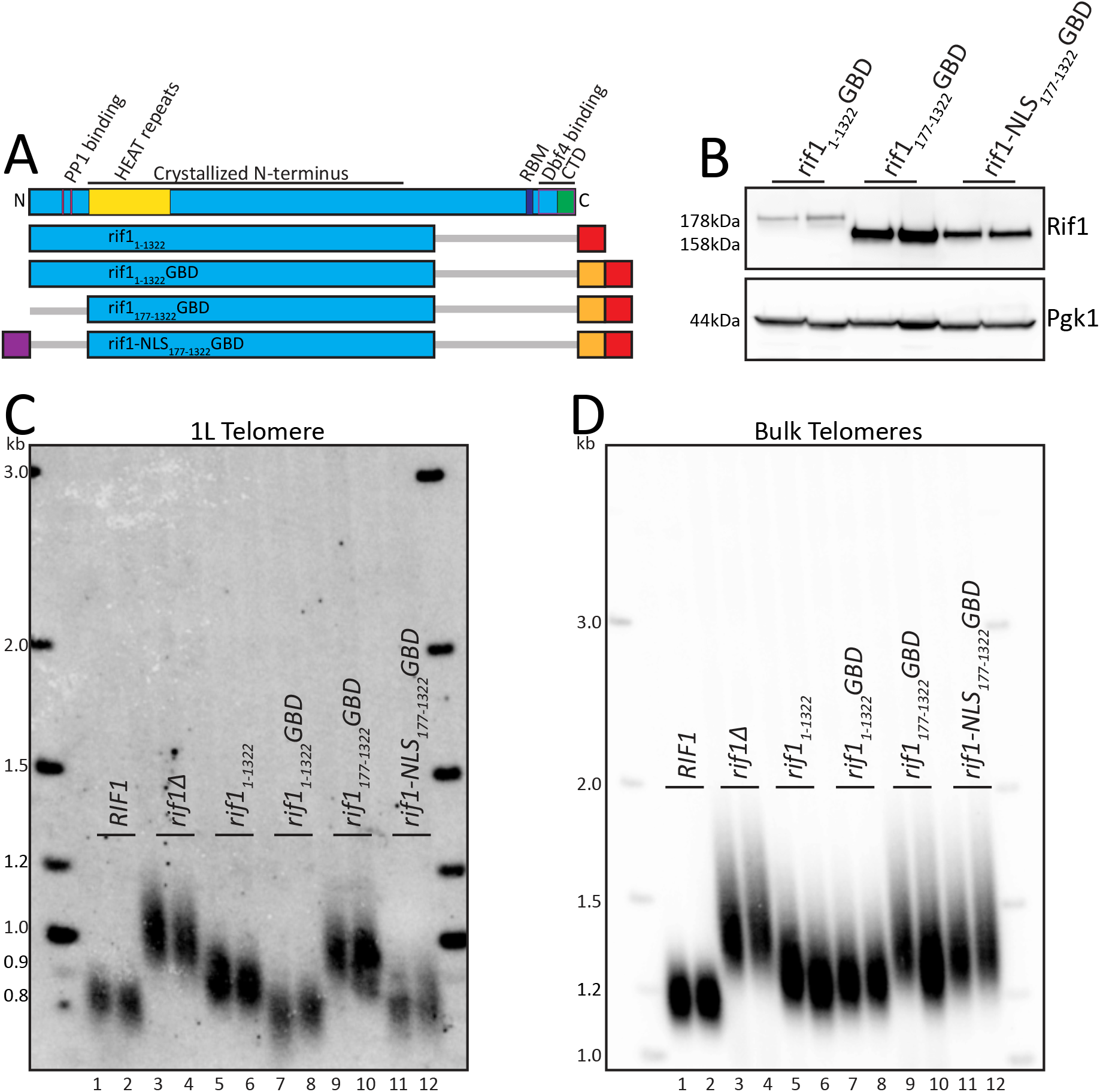
N-terminus contains critical NLS. A. RIF1 domain map as in Figure 1A. Schematic of RIF1 constructs tested in this figure (Blue: RIF1; Red: 6xFLAG; Orange: GBD; Purple: c-myc NLS; Grey bar: N- or C-terminal truncation of residues). B. Western blot showing RIF1 (anti-FLAG antibody) and Pgk1 (anti-Pgk1 antibody, control) protein levels of indicated strains. C. Southern blot showing 1L telomere probe for the indicated strains. D. Southern blot from C, rehybridized to show bulk Y’ telomere probe for the indicated strains.

### The HEAT repeats from aa 177-996 are sufficient to maintain Rif1 telomere length function

To further refine the functional region for length regulation, we made an additional truncation at the C-terminus. Truncation of Rif1 to residue 996 was previously shown to retain origin firing regulation activity (7). To test whether this truncation also retained telomere length regulation, we generated *Rif1_1-996_GBD* and *rif1-NLS_177-996_GBD* (Figure 3A), both of which were stably expressed (Figure 3B). Both *rif1_1-996_GBD* and *rif1-NLS_177-996_GBD* were able to maintain 1L telomeres at a length similar to wild type (Figure 3C, lanes 9-10 and 11-12 compared to 1-2 and 15-16). The smallest construct tested, *rif1-NLS_177-996_GBD*, was sufficient to restore nearly wild type 1L telomere length to cells that completely lack *RIF1* (Figure 3D). However, while *Rif1-NLS_177-996_GBD* was sufficient to function when tethered to the telomere, bulk telomeres remained long (Figure 3E, lanes 11-12). Rif1_177-996_ contains the HOOK domain residues 185-874 defined in Mattarocci *et al.* (14). This domain is largely comprised of HEAT repeats, a helical structural motif that can promote protein-protein interactions (23) (Figure 3A). We conclude that the HEAT repeats of Rif1 residues 177-996 are sufficient for telomere length regulation when localized to the nucleus and tethered to the telomere.

**Figure 3:**
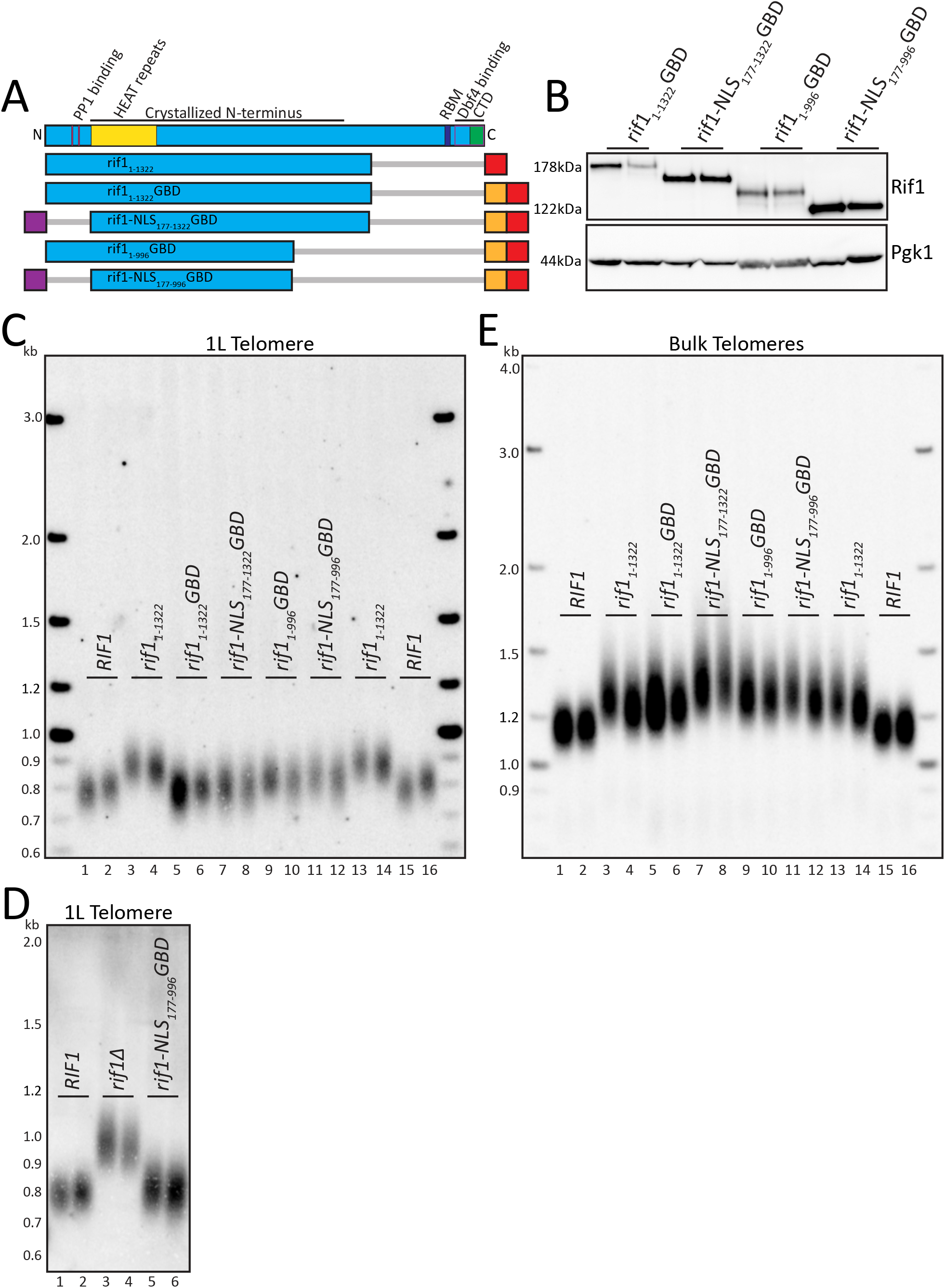
HEAT repeats of Rif1 when localized to telomere are functionally sufficient in telomere regulation. A. Rif1 domain map as in Figure 1A. Schematic of Rif1 constructs tested in this figure (Blue: Rif1; Red: 6xFLAG; Orange: GBD; Purple: c-myc NLS; Grey bar: N- or C-terminal truncation of residues). B. Western blot showing Rif1 (anti-FLAG antibody) and Pgk1 (anti-Pgk1 antibody, control) protein levels of indicated strains. C. Southern blot showing 1L telomere probe for the indicated strains. D. Southern blot showing 1L telomere probe for the indicated strains. E. Southern blot from C, rehybridized to show bulk Y’ telomere probe for the indicated strains.

### Tel1 phosphorylation is not required for HEAT repeat regulation of telomere length

Rif1 is a known substrate of Tel1 kinase (24–26), and genetic analysis indicates that *TEL1* and *RIF1* are in the same pathway of telomere length regulation (26,27). A recent paper implicated six S/T-Q motifs in the N-terminus of Rif1 as possibly playing a role in telomere length regulation (25). Four of these sites are in the Rif1_177-996_ construct and are the only S/T-Q motifs in this region. To test importance of these sites in *rif1-NLS_177-996_GBD* function, we mutated all four S/T-Q motifs (T504, S584, T775, and S824) to either alanine (A), or to a phosphomimic glutamic acid (E) (Figure 4A,B). If the role of Tel1 in telomere length acts through phosphorylation of Rif1, we predict two outcomes for telomere length: first, the A mutant should mimic short *tel1Δ* telomeres (Figure 4C, lanes 7-8) (28); and second, the E mutant should mimic long *rif1Δ* telomeres (Figure 4C, lanes 5-6). We found that that both the A and E mutants had similar telomere length to their wild type counterpart (Figure 4C, lanes 9-10, 13-14, 17-18), suggesting that phosphorylation of these S/T-Q motifs does not play a major role in length regulation. However, we note that the A mutant had slightly shorter and the E mutant had slightly longer telomeres compared to wild type, so we cannot exclude that Tel1 phosphorylation of these sites plays some minor role in Rif1 telomere regulation. Bulk telomeres also showed a subtle increase when comparing the A mutation to the E mutation in *rif1-NLS_177-996_GBD* (Figure 4D compare lanes 13-14 to 17-18). To test whether these small effects required the presence of Tel1, we made double mutants of *tel1Δ* together with *rif1-NLS_177-996 (STQ/A/E)_GBD* mutants. All double mutants had short telomeres, showing that TEL1 is epistatic to Rif1_177-996_ (Figure 4C,D, lanes 11-12, 15-16, 19-20). We conclude that Tel1 phosphorylation of Rif1 does not play a major role in Rif1 telomere regulation.

**Figure 4:**
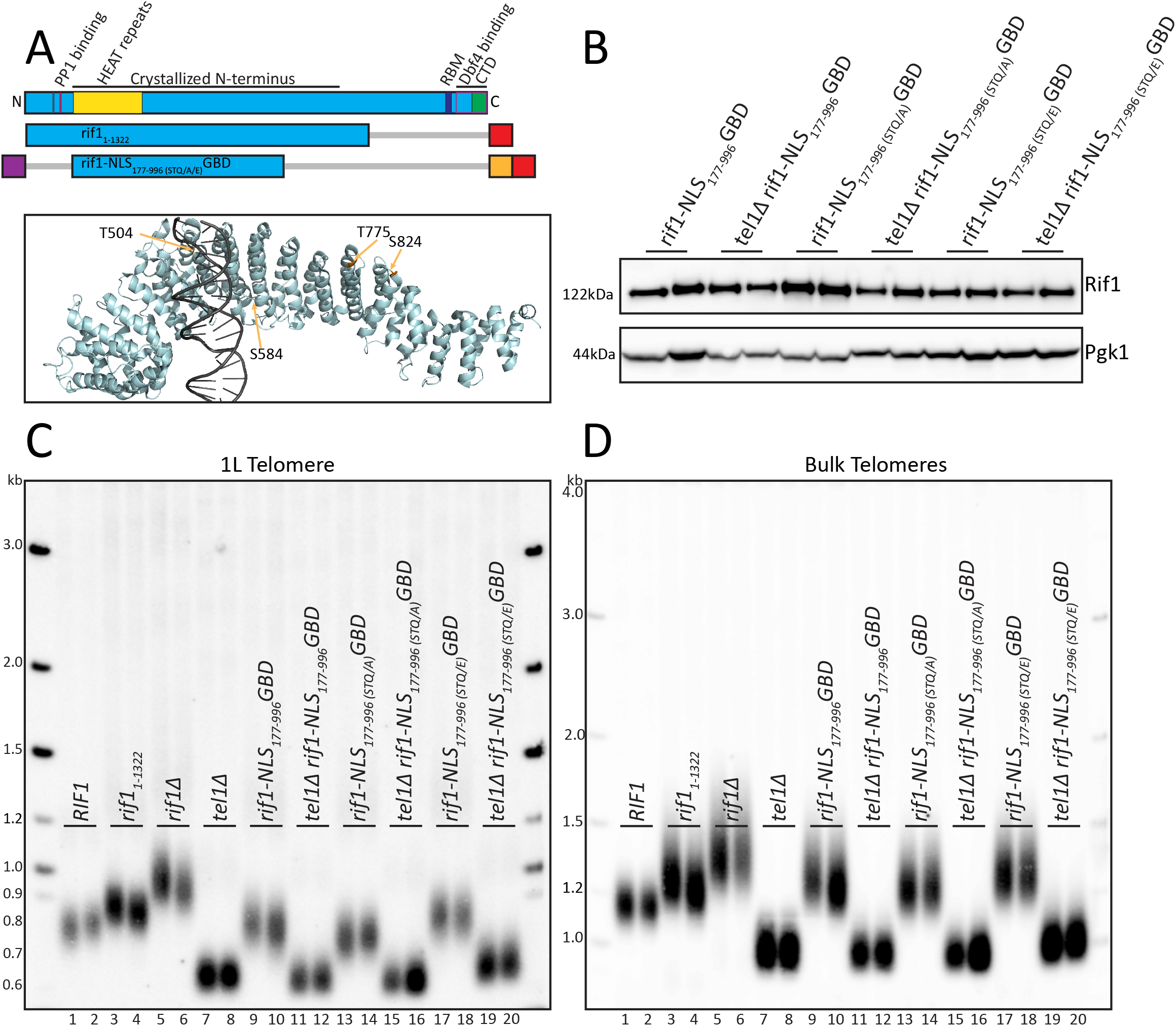
S/T-Q alanine and glutamic acid mutations have little effect on telomere length. A. Rif1 domain map as in Figure 1A. Schematic of constructs tested in this figure (Blue: Rif1; Red: 6xFLAG; Orange: GBD; Purple: c-myc NLS; Grey bar: N- or C-terminal truncation of residues). PyMOL rendering of Rif1 structure with S/T-Q residues depicted in Gold (PDB: 5NW5, showing one Rif1 monomer and DNA). B. Western blot showing Rif1 (anti-FLAG antibody) and Pgk1 (anti-Pgk1 antibody, control) protein levels of indicated strains. C. Southern blot showing 1L telomere probe for the indicated strains. D. Southern blot from C, rehybridized to show bulk Y’ telomere probe for the indicated strains.

### Positively charged residues in the HEAT repeats are required for telomere length maintenance even when tethered to the telomere

Mutations that introduce negative charges in the HEAT repeats, termed HOOK mutants, were previously shown to affect telomere length (12,14). Mattarocci *et al.* suggested that this domain mediates nonspecific binding of Rif1 to DNA via positively charged residues. Based on a crystal structure, the authors proposed that three positively charged lysine residues, K437, K563, and K570, mediate binding to the negative backbone of DNA, and they showed that a charge swap mutation to glutamic acid resulted in long telomeres (14).

To test whether the positively charged lysine residues might function to localize Rif1 to or near the telomere, or whether they might have other roles in telomere regulation, we mutated K437, K563, and K570 in our Rif1 constructs. If these lysines are mainly important for Rif1 localization to DNA, they should be dispensable at the unique 1L telomere as the GBD is sufficient to localize Rif1 to the 1L telomere. We mutated the three lysine residues, K437, K563, and K570, in both *rif1_1-1322_GBD* and *rif1-NLS_177-996_GBD*, to glutamic acid to make the same changes as in the earlier study (Figure 5A). These constructs were stably expressed (Figure 5B), but the 1L telomere length was long (Figure 5C), indicating that these mutations disrupt Rif1 telomere length function, even though the fusion protein is localized to the telomere with GBD. To further test whether the positive charge of these lysines was important, we mutated each of them to arginine, another positively charged residue, which cannot be post-translationally modified as lysine can be. We found that these arginine substitutions in *rif1-NLS_177-996 (K437R K563RK570R)_GBD* allowed Rif1 regulation of 1L telomere length (Figure 5C), demonstrating that the charge of the residues, and not their possible modification, is important for Rif1 telomere function. Finally, we mutated the three lysine residues to alanine to test whether neutral charge would allow for Rif1 telomere length function. We found that 1L telomeres were long in the alanine substitution, *rif1-NLS_177-996 (K437A K563A K570A)_GBD* (Figure 5C), supporting the conclusion that the positive charge of these residues is important for telomere length regulation. In all of the mutants, bulk telomeres were long as expected, since this construct does not localize to bulk telomeres (Figure 5D). Together, these results indicate that the positive charge of K437, K563, and K570 is important even when Rif1 is tethered to the 1L telomere, suggesting these residues are performing a function other than localizing Rif1 through binding DNA.

**Figure 5:**
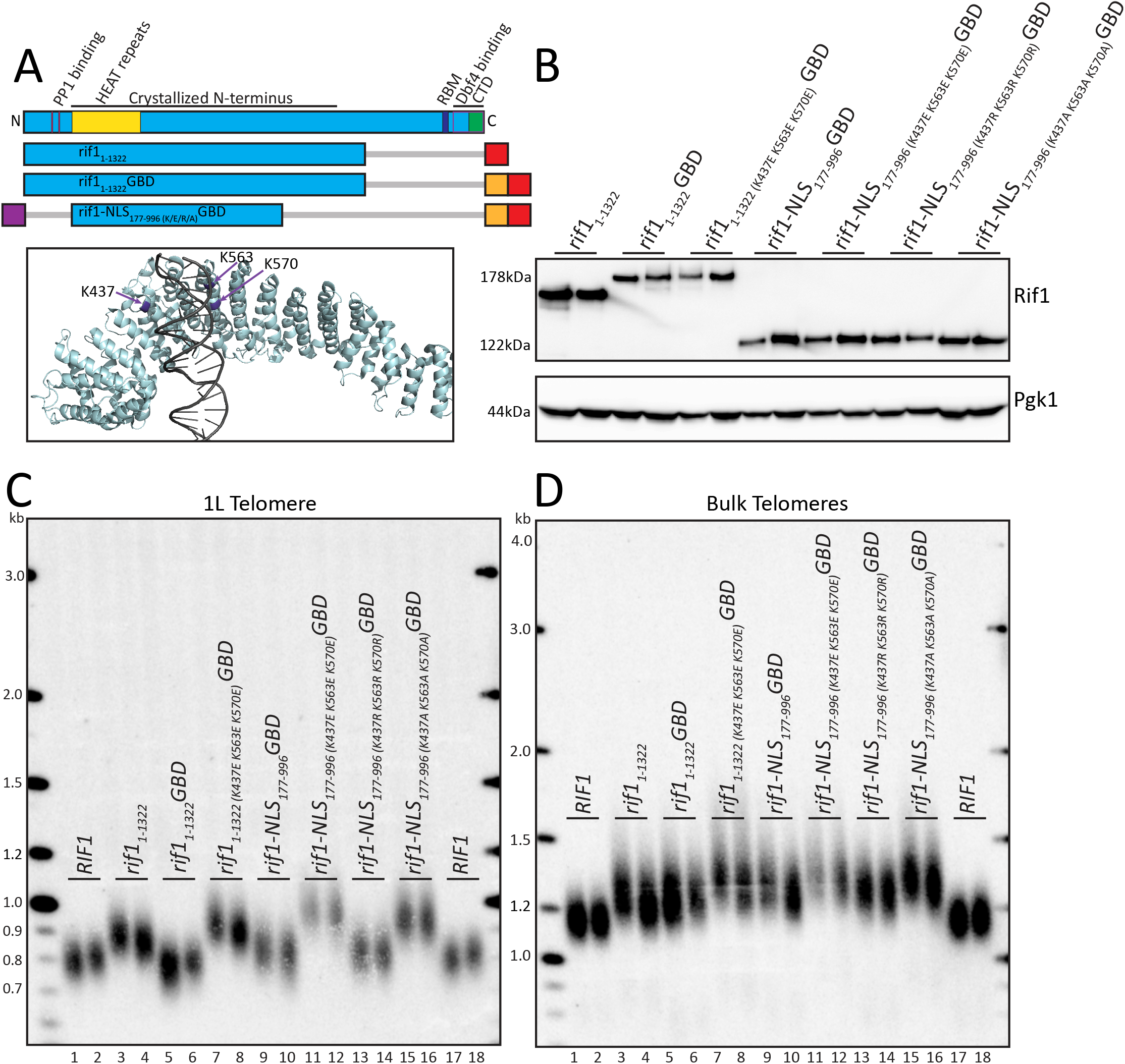
Positively charged residues in HEAT repeats are required for Rif1 function when localized to telomere. A. Rif1 domain map as in Figure 1A. Schematic of constructs tested in this figure (Blue: Rif1; Red: 6xFLAG; Orange: GBD; Purple: c-myc NLS; Grey bar: N- or C-terminal truncation of residues). PyMOL rendering of Rif1 structure with lysine residues (K437, K563, K570) depicted in Purple (PDB: 5NW5, showing one Rif1 monomer and DNA). B. Western blot showing Rif1 (anti-FLAG antibody) and Pgk1 (anti-Pgk1 antibody, control) protein levels of indicated strains. C. Southern blot showing 1L telomere probe for the indicated strains. D. Southern blot from C, rehybridized to show bulk Y’ telomere probe for the indicated strains.

### Conserved, non-positively charged, residues in HEAT repeats regulate telomere length

To further define the role of the Rif1 HEAT repeats in telomere function, we mutated conserved residues in the *rif1-NLS_177-996_GBD* construct to determine if other residues also contribute to telomere function. Because Rif1 telomere function is conserved across many yeasts (29,30), conserved residues may define regions of Rif1 needed for telomere function. First, we performed a structure-guided sequence alignment of Rif1 proteins comparing 12 yeast species (Materials and Methods; Supplementary Figure S1A,B) to identify highly conserved residues (Supplementary File 1). We found several regions of conservation throughout Rif1, similar to previous reports (13,14). These regions include the PP1 binding site, and the C-terminal Rap1-and Dbf4-binding motifs (Supplementary Figure S1C,D). The NLS that we identified is also highly conserved (Supplementary Figure S1C). Strikingly, the largest region of high conservation was in the HEAT repeats, which is included within *rif1-NLS_177-996_GBD* (Supplementary Figure S1C).

To further characterize the role of the HEAT repeats, we initially tested whether other positively charged residues in this region were functionally important. We identified conserved residues near the lysine residue cluster described above (K437, K563, and K570), which were predicted not to interact with the DNA based on the crystal structure (PDB: 5NW5). We mutated these conserved residues to alanine in three groups (Figure 6A, lanes 9-14). The PyMOL rendering of the mutated conserved residues (Gold) illustrates their relative proximity to the previously described lysines, K437, K563, and K570 (Purple), and distance from the DNA (Grey) (Figure 6C,D,E). We first tested a mutant of four residues, M436A, T564A, R565A, and W572A, each residue near one of the previously tested lysines, and found this mutant had long telomeres at the 1L telomere (Figure 6A compare lanes 9-10 to 17-18; Figure 6C; Supplementary Figure S2A). We next mutated two residues, W542A and Y532A, which are also near the DNA but not predicted to interact with it, and also found long 1L telomeres at the 1L telomere (Figure 6A compare lanes 11-12 to 17-8; Figure 6D; Supplementary Figure S2A). Finally, we mutated a conserved tyrosine, Y577, which is also predicted not to interact with DNA, and again found long telomeres at the 1L telomere (Figure 6A compare lanes 13-14 to 17-18; Figure 6E; Supplementary Figure S2A). It was striking that mutation of a single residue, Y577, had a major effect on telomere length. To further test whether Y577 is important based on its structure or its possible modification, we mutated the residue to arginine, and found that Y577R had long 1L telomeres, indicating that non-positively charged residues are also functionally important in this region of the protein (Figure 6G; Supplementary Figure S2B). This data suggests the Rif1 conserved HEAT repeats may provide as a protein binding surface, consistent with the function of other HEAT repeat domains (23).

**Figure 6:**
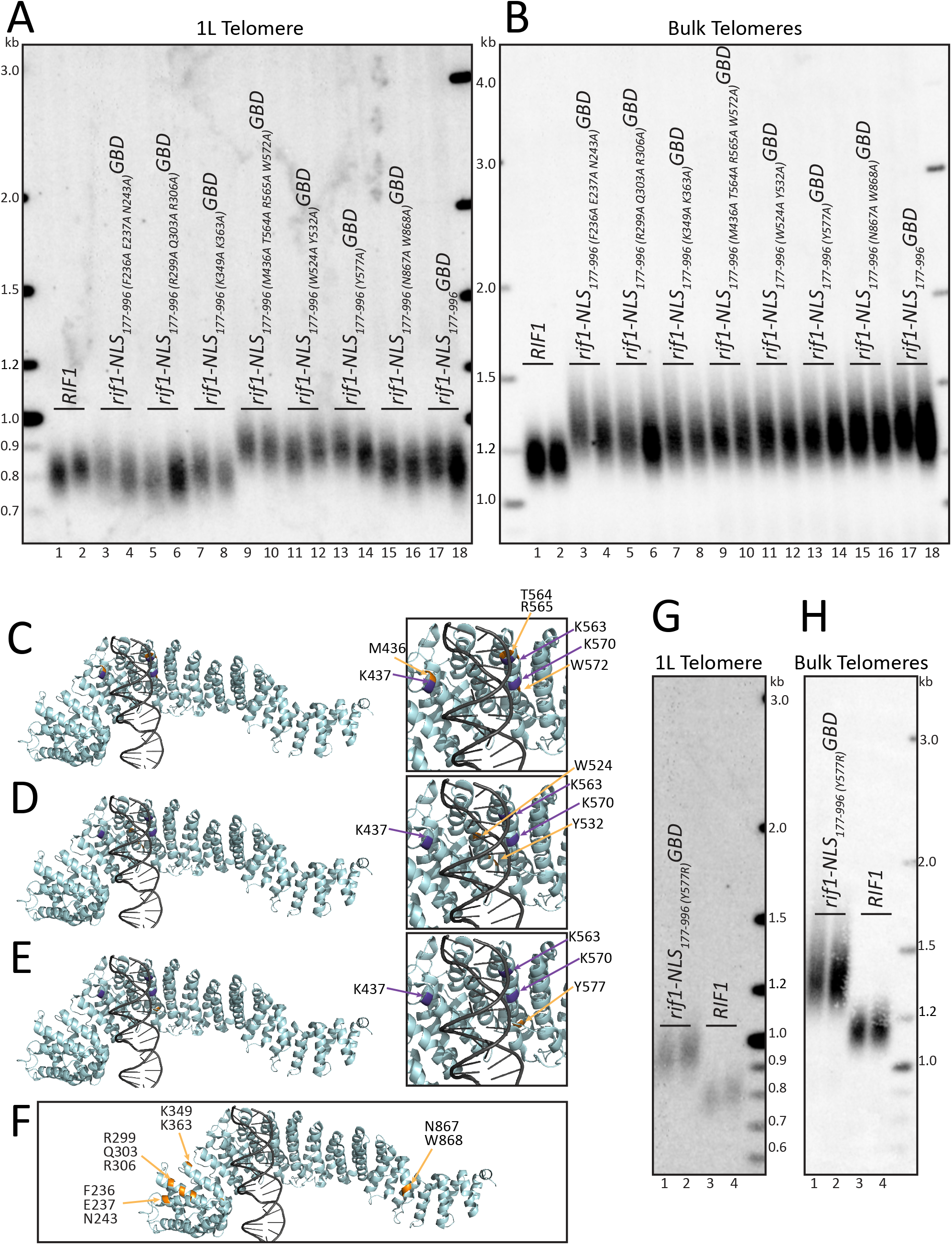
Conserved residues in HEAT repeats are critical for Rif1 function when localized to 1L telomere. A. Southern blot showing 1L telomere probe for the indicated strains. B. Southern blot from A, rehybridized to show bulk Y’ telomere probe for the indicated strains. C. PyMOL renderings of Rif1 structure. Purple: lysines (K437, K563, K570). Gold: residues mutated in strains from lanes 9-10 in above Southern blots (M436, T564, R565, W572) (PDB: 5NW5, showing one Rif1 monomer and DNA). D. PyMOL renderings of Rif1 structure. Purple: lysines (K437, K563, K570). Gold: residues mutated in strains from lanes 11-12 in above Southern blots (W524, Y532) (PDB: 5NW5, showing one Rif1 monomer and DNA). E. PyMOL renderings of Rif1 structure. Purple: lysines (K437, K563, K570). Gold: residues mutated in strains from lanes 13-14 in above Southern blots (Y577) (PDB: 5NW5, showing one Rif1 monomer and DNA). F. PyMOL rendering of Rif1 structure. Purple: lysines (K437, K563, K570). Gold: residues mutated in strains from lanes 3-8 and 15-16 in above Southern blots (PDB: 5NW5, showing one Rif1 monomer and DNA). G. Southern blot showing 1L telomere probe of the indicated strains. H. Southern blot from F, rehybridized to show bulk Y’ telomere probe for the indicated strains.

When we mutated highly conserved residues that were located distal to the previously characterized lysines (K437, K563, K570), we found these mutations did not affect telomere length (Figure 6F; Supplementary Figure S2A). A first mutant containing three residue substitutions, F236A, E237A, and N242A, a second mutant containing three other substitutions, R299A, Q303A, and R306A, and a third mutant with two residue substitutions, K349A and K363A, all had a 1L telomere length similar to the *rif1-NLS_177-996_GBD* control construct (Figure 6A compare lanes 3-4, 5-6, 7-8 to 1-2; Figure 6F), indicating they do not disrupt Rif1 function. We saw some clonal variation in both the (F236A, E237A, N242A) and (R299A Q303A, R306A) mutants, but found the majority of these mutants showed telomere length similar to the *rif1-NLS_177-996_GBD* control. Our data suggest that conserved residues distal from the K437, K563, and K570 lysine residues, regardless of charge, are not important in Rif1 telomere length regulation.

The crystal structure suggests that central residues within Rif1 might mediate a dimer interface in the presence of DNA (14). To test the function of this putative dimer interface on telomere regulation, we mutated two residues, N867 and W868, which are the most highly conserved residues in the region (Figure 6F; Supplementary Figure S2A). We found that this mutant did not disrupt Rif1 function, as 1L telomere length was similar to that of the *Rif1-NLS_177-996_GBD* control (Figure 6A compare lanes 15-16 to 17-18; Figure 6F). This result suggests that Rif1 dimerization may not be critical for function in telomere length regulation. The bulk telomeres of all *rif1-NLS_177-996_GBD* mutants tested were long, as expected (Figure 6B,H). We have summarized the telomere lengths of the different mutants in Supplementary Figure S2C. Our data demonstrates that the conserved residues required for Rif1 telomere function cluster around the positively charged lysine residues, K437, K563, K570. Our data do not support a model for the HEAT repeats simply mediating Rif1 DNA binding and potential telomere localization, since the lysine residues are important even when bound to the telomere, and both charged and uncharged residues compromise Rif1 function. Instead, we propose that this region may mediate a protein-protein interaction required for telomere length regulation.

## DISCUSSION

Rif1 has been known to regulate telomere length for over two decades, but we do not yet understand the mechanism of this regulation. We took a mutational approach to determine which regions of Rif1 are necessary for Rif1 function at the telomere. We localized different portions of Rif1 fused to the Gal4 DNA binding domain to a unique telomere at 1L. This approach enabled us to compare 1L telomere length to bulk telomere length to probe regions of Rif1 that function at the telomere. We identified a functional domain of Rif1, which, when affixed with an NLS, was sufficient to maintain telomere length similar to wild type length at the unique telomere. This conserved region of HEAT repeats from aa 177-996 is comprised both of positively charged residues as well as other critical polar and nonpolar conserved residues, which are required for telomere length regulation. Our data indicate that the Rif1 conserved HEAT repeats may provide a surface for protein-protein interactions that optimally functions in telomere length regulation when more Rif1 is at the telomere.

### Conserved NLS required for localization of Rif1 domains

We identified a highly conserved NLS in the Rif1 NTD, which is critical for telomere length regulation. Indeed, the functional role of the NTD can be replaced by an exogenous NLS and wild type telomere length was restored. This deletion analysis demonstrates that the primary role of the NTD is to localize Rif1 to the nucleus, and that the PP1 binding site is not required for telomere length regulation. This finding supports our previous mutational analysis showing that substitutions in the PP1 binding region of Rif1 do not affect telomere length regulation (12). Identification of the NLS in the N-terminus is also of interest because in fruit flies and vertebrates a conserved NLS is present in the C-terminus (31,32). While Rif1 is conserved from yeast through humans, many functional domains of Rif1 appear to have been rearranged throughout evolution. For example, the N-terminal PP1 binding site is located in the C-terminus in mammalian Rif1 (15). Conservation of these regions suggests that, while there has been some rearrangement of domains, there may be evolutionary pressure to maintain these functional domains in Rif1 homologues.

### Rif1 HEAT repeats region is sufficient to regulate telomere length when localized to telomeres

The Rif1 region from aa 177-996, when tethered to the telomere, was able to maintain 1L telomeres similar to wild type length. This region consists of conserved HEAT repeats, which typically promote protein-protein interactions (15,23). These HEAT repeats function independently of the most N-terminal region of aa 1-176 and the C-terminus of aa 997-1916 to maintain telomere length when bound to the telomere. This indicates that known functions in the N-and C-termini, namely PP1 binding, Dbf4 binding, and Rap1 binding, are not required for Rif1 telomere function when Rif1 is tethered to the telomere.

The Rif1 HEAT repeats likely mediate protein-protein interaction, not just DNA binding. It was recently shown that Rif1 crystalizes with DNA (14), and the authors proposed that this DNA binding may add an additional mechanism to localize Rif1 to telomeres. Of note, Rif1 also crystallized as a monomer without DNA, in addition to crystallizing as a dimer with DNA (PDB: 5NVR and 5NW5, respectively) (14). Here we confirmed that the lysine residues implicated in DNA binding have functional importance in regulating telomere length, but their function likely extends beyond DNA binding, as they are required for telomere length regulation even when tethered to the telomere (Figure 5). Moreover, several conserved residues proximal to these lysine residues, which were not predicted to interact with DNA, were also critical for telomere length regulation. However, distal conserved residues did not affect telomere length (Figure 6C,D,E vs Figure 6F). The small surface around these lysine residues (K437, K563, K570), between M436 and Y577, which has the greatest impact on telomere function, is also the most conserved HEAT repeat from yeast to humans (15). This conserved region, therefore, seems to function in telomere length regulation beyond localizing Rif1 to DNA.

### Rif1 HEAT repeats have partial function when not tethered to the telomere

While Rif1 HEAT repeat constructs restored wild type telomere length when localized by GBD to 1L, we found that they also had a small effect on bulk telomere length (Figure 1D, Figure 3E, Figure 4D). This was surprising as Rif1_1-1322_ and Rif1-NLS_177-996_ both lack Rap1 binding and, therefore, cannot localize to the telomere (13,33,34). For Rif1_1-1322_, this partial effect has been previously reported (12,13). These data suggest that these constructs can function in telomere length regulation by an unknown mechanism, which is independent of Rap1 binding. We suggest that localizing Rif1 to the telomere, through either Rap1 binding or GBD, promotes a high local concentration of Rif1; however, Rif1 can still partially perform its function of regulating telomere length when not telomere bound. Perhaps there is simply a lower local concentration when Rif1 is not tethered to the telomere, leading to the partial length regulation that is seen. This conclusion is supported by longstanding evidence that recruiting more Rif1 to the telomere, by adding Rap1 binding sites or by directly tethering full length Rif1, leads to progressively shorter telomeres (35,36).

In spite of a high degree of conservation of Rif1 homologues in origin firing, its role as a telomere regulator has only been characterized in yeasts (37–39). Moreover, localization to telomeres has only been shown in yeast, although through different mechanisms (29). We have shown that even heterologous tethering of Rif1 to a telomere negatively regulates telomere length. This suggests the possibility that, if Rif1 were it present at higher concentration at telomeres in other organisms, it might exert a negative effect on telomere elongation.

### Evolutionarily conserved HEAT repeats are important in both NHEJ and telomere length

The Rif1 HEAT repeats region, which we found is critical for carrying out telomere length regulation, is also implicated in promoting NHEJ in both yeast and mammals. This region, which contains the positively charged lysines (K437, K563, K570), is required to promote NHEJ in yeast, and this effect does not require binding to Rap1 (14). The importance of the positively charged residues in the HEAT repeats may indicate that Rif1 is binding to a negatively charged protein domain or, possibly, a phosphorylated protein. In mammalian cells, Rif1 HEAT repeats are specifically required for Rif1 localization at double-strand breaks (18). The ATM-dependent phosphorylation of 53BP1 recruits Rif1 to sites of DNA damage to block end resection and promote NHEJ (16–19,39,40). This suggests a possible role for the 53BP1 yeast ortholog Rad9, which is similarly phosphorylated at S/T-Q residues by Tel1/Mec1 (41,42). However, Rad9 does not have a telomere length effect on its own, so perhaps the HEAT repeats bind to and regulate the activity of another protein to promote both NHEJ and telomere length regulation. Additionally, while Tel1 functions in the same genetic pathway as Rif1 (26), its role in telomere length regulation is also not fully understood. Our data indicate that Tel1 phosphorylation of Rif1 is not the main mechanism by which Tel1 regulates telomere length (Figure 4). Understanding the roles of Tel1 phosphorylation and Rif1 in both telomere length regulation and NHEJ will allow more complete understanding of Rif1 function.

We have dissected the domains of Rif1 that function to regulate telomere length. We found that the Rif1 HEAT repeats region between M436 and Y577 is primarily responsible for Rif1 telomere function, and that the N-and C-termini of Rif1 are not required for Rif1 function in telomere length regulation. While we cannot rule out some role for DNA binding of this region, our data is most consistent with the Rif1 HEAT repeats interacting with a protein partner to carry out its telomere function. The partial function of Rif1 when not bound to the telomere suggests that the local concentration of Rif1 is important for its function in telomere length regulation. These results suggest several exciting questions to understand Rif1 function in telomere length regulation: are these functions conserved or yeast specific?, which protein partners are interacting with Rif1 to carry out these functions?, and is the conserved role of Rif1 in NHEJ commandeered in yeast for telomere regulation?

## ACKNOWLEDGEMENT

We thank Dr. Brendan Cormack and Dr. Thomas Kelly for insightful discussions in planning experiments and discussing data. We would like to acknowledge Dr. Brendan Cormack, Dr. Thomas Kelly, Dr. Rebecca Keener, Sam Sholes, Margaret Strong, Carla Connelly and Dr. Akshi Jasani for critical reading of the manuscript.

---

**Supplementary Figure S1:**
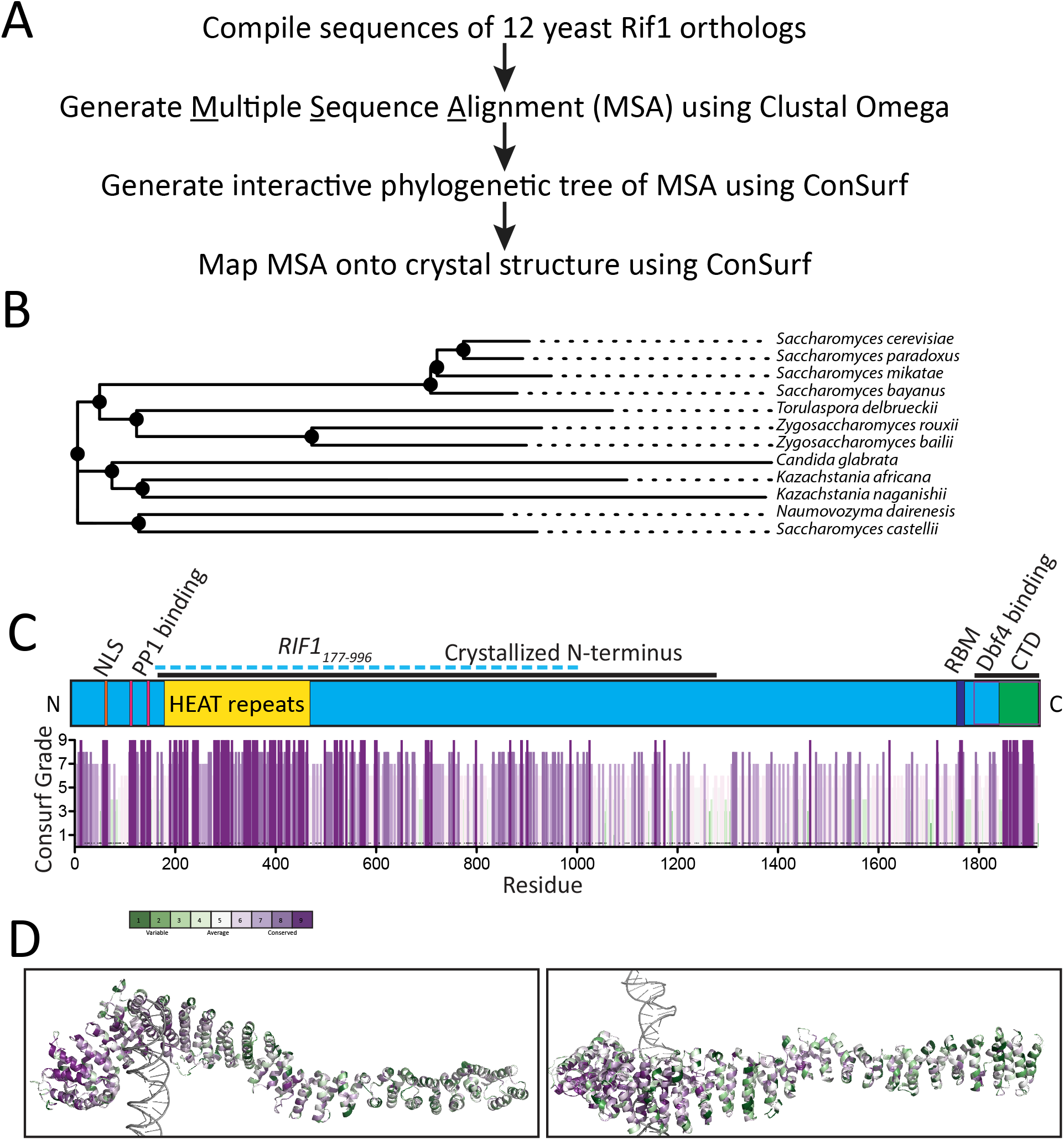
Structure-based sequence alignment. A. Workflow of structure-based sequence alignment (see Materials and Methods). B. Phylogenetic tree of the Multiple Sequence Alignment (MSA) generated using ConSurf. C. Conservation graph generated using GraphPad Prism of sequence alignment with position in Rif1 protein on the X-axis. The height of the bar indicates ConSurf Grade, with a higher bar having higher conservation. Purple indicates high conservation, and green indicates low conservation. Rif1 domain map is depicted to scale above the bar graph, with the newly defined NLS added, and Rif1 aa 177-196 are depicted with a blue dotted line (RBM: Rap1 binding motif; CTD: Carboxyl-terminal domain; Dbf4 binding overlaps CTD). D. MSA mapped onto the Rif1 crystal structure (PDB: 5NW5, showing one Rif1 monomer and DNA) visualized using PyMOL. Purple indicates high conservation, and green indicates low conservation.

**Supplementary Figure S2:**
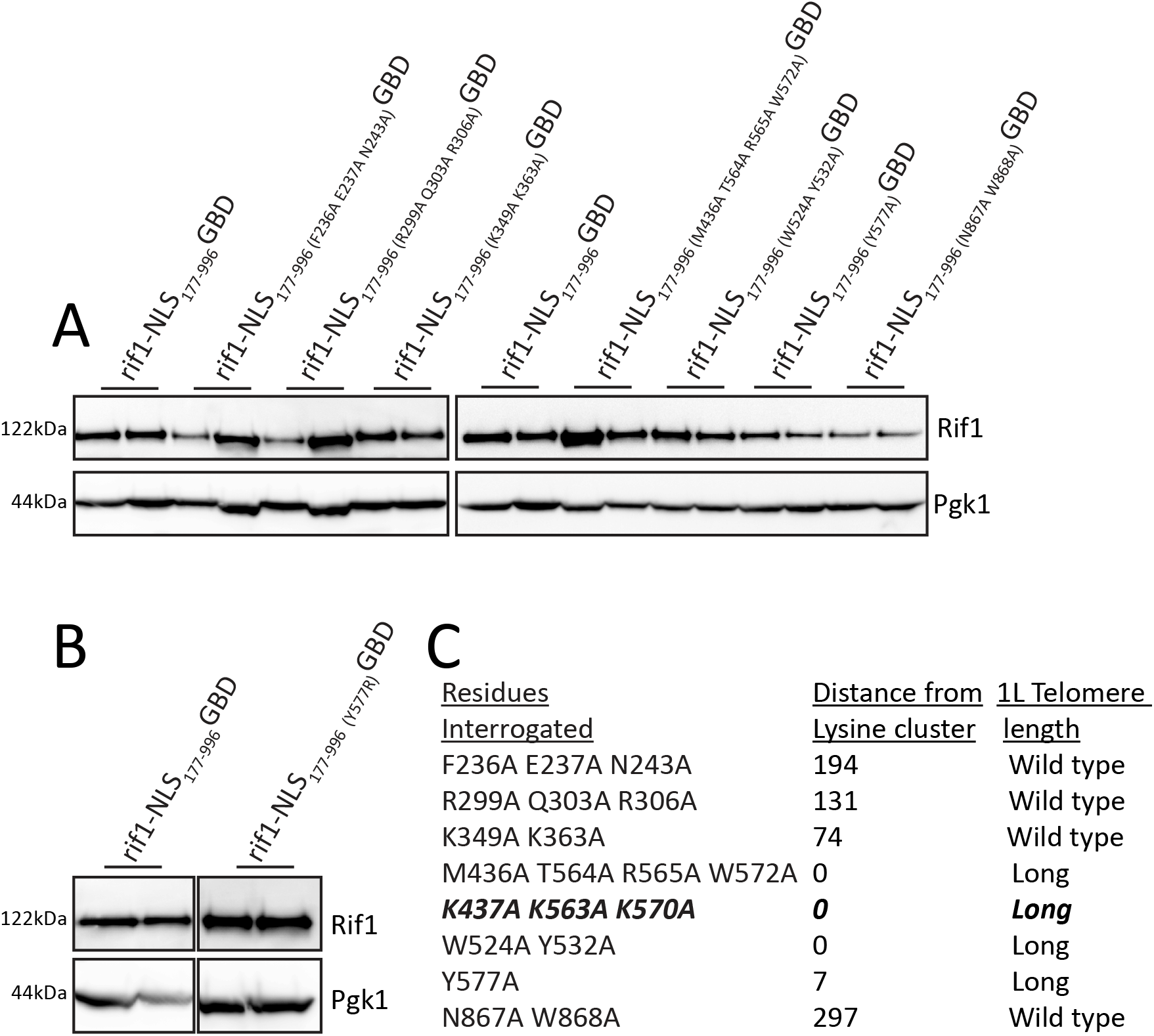
Protein levels for strains mutated in conserved HEAT repeats. A. Western blot showing Rif1 (anti-FLAG antibody) and Pgk1 (anti-Pgk1 antibody, control) protein levels of indicated strains, related to Figure 6A,B. B. Western blot showing Rif1 (anti-FLAG antibody) and Pgk1 (anti-Pgk1 antibody, control) protein levels of indicated strains, related to Figure 6F,G. C. Conserved residues with corresponding distance from lysine cluster and 1L telomere length, related to Figure 6A.

**Supplementary Table 1: Strain list, plasmid list, and cloning specifics and reagents**

**Supplementary File 1: Multiple Sequence Alignment (MSA) of Rif1**

MSA of Rif1 generated as described in Materials and Methods and visualized using SnapGene software.

## REFERENCES

1. Palm, W. and de Lange, T. (2008) How shelterin protects mammalian telomeres. Annu Rev Genet, 42, 301–334.

2. Wellinger, R.J. and Zakian, V.A. (2012) Everything you ever wanted to know about Saccharomyces cerevisiae telomeres: beginning to end. Genetics, 191, 1073–1105.

3. Harley, C.B., Futcher, A.B. and Greider, C.W. (1990) Telomeres shorten during ageing of human fibroblasts. Nature, 345, 458–460.

4. Armanios, M. and Blackburn, E.H. (2012) The telomere syndromes. Nat Rev Genet, 13, 693–704.

5. McNally, E.J., Luncsford, P.J. and Armanios, M. (2019) Long telomeres and cancer risk: the price of cellular immortality. J Clin Invest, 130, 3474–3481.

6. Hardy, C.F., Sussel, L. and Shore, D. (1992) A RAP1-interacting protein involved in transcriptional silencing and telomere length regulation. Genes Dev, 6, 801–814.

7. Hiraga, S., Alvino, G.M., Chang, F., Lian, H.Y., Sridhar, A., Kubota, T., Brewer, B.J., Weinreich, M., Raghuraman, M.K. and Donaldson, A.D. (2014) Rif1 controls DNA replication by directing Protein Phosphatase 1 to reverse Cdc7-mediated phosphorylation of the MCM complex. Genes Dev, 28, 372–383.

8. Mattarocci, S., Shyian, M., Lemmens, L., Damay, P., Altintas, D.M., Shi, T., Bartholomew, C.R., Thoma, N.H., Hardy, C.F. and Shore, D. (2014) Rif1 controls DNA replication timing in yeast through the PP1 phosphatase Glc7. Cell reports, 7, 62–69.

9. Dave, A., Cooley, C., Garg, M. and Bianchi, A. (2014) Protein phosphatase 1 recruitment by Rif1 regulates DNA replication origin firing by counteracting DDK activity. Cell reports, 7, 53–61.

10. Hiraga, S.I., Ly, T., Garzon, J., Horejsi, Z., Ohkubo, Y.N., Endo, A., Obuse, C., Boulton, S.J., Lamond, A.I. and Donaldson, A.D. (2017) Human RIF1 and protein phosphatase 1 stimulate DNA replication origin licensing but suppress origin activation. EMBO reports, 18, 403–419.

11. Alver, R.C., Chadha, G.S., Gillespie, P.J. and Blow, J.J. (2017) Reversal of DDK-Mediated MCM Phosphorylation by Rif1-PP1 Regulates Replication Initiation and Replisome Stability Independently of ATR/Chk1. Cell reports, 18, 2508–2520.

12. Shubin, C.B. and Greider, C.W. (2020) The role of Rif1 in telomere length regulation is separable from its role in origin firing. eLife, 9.

13. Shi, T., Bunker, R.D., Mattarocci, S., Ribeyre, C., Faty, M., Gut, H., Scrima, A., Rass, U., Rubin, S.M., Shore, D. et al. (2013) Rif1 and Rif2 shape telomere function and architecture through multivalent Rap1 interactions. Cell, 153, 1340–1353.

14. Mattarocci, S., Reinert, J.K., Bunker, R.D., Fontana, G.A., Shi, T., Klein, D., Cavadini, S., Faty, M., Shyian, M., Hafner, L. et al. (2017) Rif1 maintains telomeres and mediates DNA repair by encasing DNA ends. Nat Struct Mol Biol, 24, 588–595.

15. Sreesankar, E., Senthilkumar, R., Bharathi, V., Mishra, R.K. and Mishra, K. (2012) Functional diversification of yeast telomere associated protein, Rif1, in higher eukaryotes. BMC genomics, 13, 255.

16. Di Virgilio, M., Callen, E., Yamane, A., Zhang, W., Jankovic, M., Gitlin, A.D., Feldhahn, N., Resch, W., Oliveira, T.Y., Chait, B.T. et al. (2013) Rif1 prevents resection of DNA breaks and promotes immunoglobulin class switching. Science, 339, 711–715.

17. Zimmermann, M., Lottersberger, F., Buonomo, S.B., Sfeir, A. and de Lange, T. (2013) 53BP1 regulates DSB repair using Rif1 to control 5’ end resection. Science, 339, 700–704.

18. Escribano-Diaz, C., Orthwein, A., Fradet-Turcotte, A., Xing, M., Young, J.T., Tkac, J., Cook, M.A., Rosebrock, A.P., Munro, M., Canny, M.D. et al. (2013) A cell cycle-dependent regulatory circuit composed of 53BP1-RIF1 and BRCA1-CtIP controls DNA repair pathway choice. Mol Cell, 49, 872–883.

19. Chapman, J.R., Barral, P., Vannier, J.B., Borel, V., Steger, M., Tomas-Loba, A., Sartori, A.A., Adams, I.R., Batista, F.D. and Boulton, S.J. (2013) RIF1 is essential for 53BP1-dependent nonhomologous end joining and suppression of DNA double-strand break resection. Mol Cell, 49, 858–871.

20. Mitchell, L.A. and Boeke, J.D. (2014) Circular permutation of a synthetic eukaryotic chromosome with the telomerator. Proc Natl Acad Sci U S A, 111, 17003–17010.

21. Gallo, C.M., Smith, D.L., Jr. and Smith, J.S. (2004) Nicotinamide clearance by Pnc1 directly regulates Sir2-mediated silencing and longevity. Mol Cell Biol, 24, 1301–1312.

22. Dang, C.V. and Lee, W.M. (1988) Identification of the human c-myc protein nuclear translocation signal. Mol Cell Biol, 8, 4048–4054.

23. Andrade, M.A., Petosa, C., O’Donoghue, S.I., Muller, C.W. and Bork, P. (2001) Comparison of ARM and HEAT protein repeats. Journal of molecular biology, 309, 1–18.

24. Smolka, M.B., Albuquerque, C.P., Chen, S.H. and Zhou, H. (2007) Proteome-wide identification of *in vivo* targets of DNA damage checkpoint kinases. Proc Natl Acad Sci U S A, 104, 10364–10369.

25. Wang, J., Zhang, H., Al Shibar, M., Willard, B., Ray, A. and Runge, K.W. (2018) Rif1 phosphorylation site analysis in telomere length regulation and the response to damaged telomeres. DNA Repair (Amst), 65, 26–33.

26. Sridhar, A., Kedziora, S. and Donaldson, A.D. (2014) At short telomeres Tel1 directs early replication and phosphorylates Rif1. PLoS Genet, 10, e1004691.

27. Craven, R.J. and Petes, T.D. (1999) Dependence of the regulation of telomere length on the type of subtelomeric repeat in the yeast Saccharomyces cerevisiae. Genetics, 152, 1531–1541.

28. Greenwell, P.W., Kronmal, S.L., Porter, S.E., Gassenhuber, J., Obermaier, B. and Petes, T.D. (1995) TEL1, a gene involved in controlling telomere length in S. cerevisiae, is homologous to the human ataxia telangiectasia gene. Cell, 82, 823–829.

29. Kanoh, J. and Ishikawa, F. (2001) spRap1 and spRif1, recruited to telomeres by Taz1, are essential for telomere function in fission yeast. Curr Biol, 11, 1624–1630.

30. Castano, I., Pan, S.J., Zupancic, M., Hennequin, C., Dujon, B. and Cormack, B.P. (2005) Telomere length control and transcriptional regulation of subtelomeric adhesins in Candida glabrata. Mol Microbiol, 55, 1246–1258.

31. Batenburg, N.L., Walker, J.R., Noordermeer, S.M., Moatti, N., Durocher, D. and Zhu, X.D. (2017) ATM and CDK2 control chromatin remodeler CSB to inhibit RIF1 in DSB repair pathway choice. Nat Commun, 8, 1921.

32. Xu, D., Muniandy, P., Leo, E., Yin, J., Thangavel, S., Shen, X., Ii, M., Agama, K., Guo, R., Fox, D., 3rd et al. (2010) Rif1 provides a new DNA-binding interface for the Bloom syndrome complex to maintain normal replication. Embo J, 29, 3140–3155.

33. Hiraga, S.I., Monerawela, C., Katou, Y., Shaw, S., Clark, K.R., Shirahige, K. and Donaldson, A.D. (2018) Budding yeast Rif1 binds to replication origins and protects DNA at blocked replication forks. EMBO reports, 19.

34. Hafner, L., Lezaja, A., Zhang, X., Lemmens, L., Shyian, M., Albert, B., Follonier, C., Nunes, J.M., Lopes, M., Shore, D. et al. (2018) Rif1 Binding and Control of Chromosome-Internal DNA Replication Origins Is Limited by Telomere Sequestration. Cell reports, 23, 983–992.

35. Marcand, S., Gilson, E. and Shore, D. (1997) A protein-counting mechanism for telomere length regulation in yeast. Science, 275, 986–990.

36. Levy, D.L. and Blackburn, E.H. (2004) Counting of Rif1p and Rif2p on Saccharomyces cerevisiae telomeres regulates telomere length. Molecular and cellular biology, 24, 10857–10867.

37. Xu, L. and Blackburn, E.H. (2004) Human Rif1 protein binds aberrant telomeres and aligns along anaphase midzone microtubules. The Journal of cell biology, 167, 819–830.

38. Buonomo, S.B., Wu, Y., Ferguson, D. and de Lange, T. (2009) Mammalian Rif1 contributes to replication stress survival and homology-directed repair. J Cell Biol, 187, 385–398.

39. Silverman, J., Takai, H., Buonomo, S.B., Eisenhaber, F. and de Lange, T. (2004) Human Rif1, ortholog of a yeast telomeric protein, is regulated by ATM and 53BP1 and functions in the S-phase checkpoint. Genes Dev, 18, 2108–2119.

40. Feng, L., Fong, K.W., Wang, J., Wang, W. and Chen, J. (2013) RIF1 counteracts BRCA1-mediated end resection during DNA repair. J Biol Chem, 288, 11135–11143.

41. Vialard, J.E., Gilbert, C.S., Green, C.M. and Lowndes, N.F. (1998) The budding yeast Rad9 checkpoint protein is subjected to Mec1/Tel1-dependent hyperphosphorylation and interacts with Rad53 after DNA damage. Embo J, 17, 5679–5688.

42. Emili, A. (1998) MEC1-Dependent Phosphorylation of Rad9p in Response to DNA Damage. Molecular Cell, 2, 183–189.

