## Supplementary File 1 for "Rif1 regulates telomere length through conserved HEAT repeats"

### Aligned using Clustal Omega

**Consensus**
**1.** Saccharomyces_cerevisiae
**2.** Kazachstania_naganishii
**3.** Candida_glabrata
**4.** Saccharomyces_castellii
**5.** Naumovozyma_dairenensis
**6.** Kazachstania_africana
**7.** Torulaspora_delbrueckii
**8.** Saccharomyces_mikatae
**9.** Saccharomyces_bayanus
**0.** Saccharomyces_paradoxus
**1.** Zygoaccharomyces_bailii
**2.** Zygosaccharomyces_rouxii

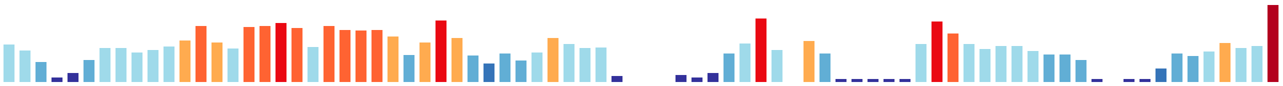

```
    MSX--XXXXXXXKXXRAIDXLDXHIXXRSXXXXXSXXX------XXXWX-XX-----XLPXXXXXXXX----XXXXXXXS

1   -----MSKDFSDKKKHTIDRIDQHILRRSQHDNYSNGS-------SPWM-KT-----NLPP-PSPQAH----MHIQSDLS   57
2   MTEHSTERSSNRMINRTLEMLEKAGSKKNT---FAN--------KYPWS-SGNFKRNNAPIPEHTD-------SRDYSTS   61
3   MSD---GSADRKGKMRALDILSKHLRGKNSDKTTPTRYGHKSVELESILRKDNIVSSPLESRDHQESQR-INNPHSNTAS   76
4   MSDTPTKNTSNKKSDRAIDLLNKQLKLK-----NSRSSP---IIPLQWF-NS-----QVSQQESS--------FNPNELS   58
5   MSDSSKESSGPRKNDRAITLLDKHINQKSKMKSTSNSSP---IVLTQWF-NS-----QLPAPPSTSPVSVQTHKAANDVS   71
6   MSDESHKDASPKKNMKALAMIDKHISQRTLQKKNLSPS-------STWV-KP-SMSNQLSAQPQQNAAY---WNSANNTS   68
7   MSTKPDGDMHREKKMRAIDMLNHFINKRSQAKAKSVSPT-----ARNWK-NS-----RLSPPNTEEPN-----EDANNVS   64
8   -----MSKDFSDKKKHTIDRIDQHILRRSQHDVNPNGS-------SPWM-KT-----KLPPLPFPQAH----MHIQSDLS   58
9   -----MSNDFSDKKKHTIDRIDQHILRRSQQDAHSSGN-------SPWM-KA-----KLPPPPSPQAH----MHMQSDLS   58
10  -----MSKDFSDKKKHTIDRIDQHILRRSQHDAHSNGS-------SPWM-KV-----NLPPPPSPQAQ----MHIRSDLS   58
11  MSS--PPGQSTEKRMRAIDRLDQHMIKRSKSNNRASSP----VPFLSWI-TQ-----KLPPPSSLLGQ----RDAPTQTS   64
12  MSS--SSNQSSEKRMRAIDRLDQHMMKRSMKNSKSSSPS---GPFLPWL-SH-----KLPSNSQSDSNG---SQNHEISS   66
```

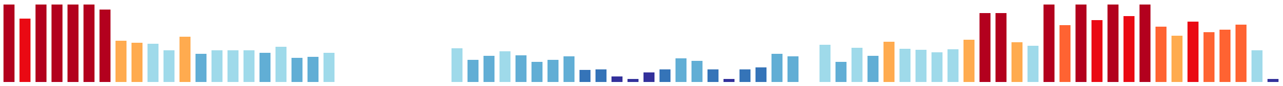

```
    PTPKRRKXXXXSXXXXXXXXX-------XXXXXXXXXX--XXXXX-XXXX-EXXXSXXXXXSPXXKSVAFSDRXESSPX-

1   PTPKRRKLASSSDCENKQFDL-------SAINKNLYPEDTGSRLM-QSLP-ELSASNSDNVSPVTKSVAFSDRIESSPI-   127
2   PTPKRRKMATPQSADKTTVQA-------NSNSLDNLK----ISL---VFP-KDSERPLSSSSPCQKSVSFSDRIEVESSS   126
3   PTPKRRKLNTQAISSDNKTKQVLSIGELSHPDMQSTPA--RSRIMKKDIGDQNMSSSEIYSPSVNKSVSFLDESNKNSE-   153
4   PTPKRRKLNNKRPATEKSFKK-------NAINLEETPILMAPKSISDTR----FDDPISNSSPIRKAVAFSDKVESSPTQ   127
5   PTPKRRRVNNMDQTSSTVPHL----------DLRRF------ELNNDTRL-ERSTSESNSASPIRKTVAFSDKIEIESSP   134
6   PTPKRRKITSSSEATLPEGSK-------NFTVLNKAFE----------VV-GQGPSEDILSSPGQKSVAFSDRIDSSPE-   129
7   PTPKRRKFAQSSPVRKPEEPV-------LSTDV-------QLK----LPA-TESLNDKAEGSPGTKSVAFSDKLDSSPT-   124
8   PTPKRRKLAFSSNYDNKELDL-------SANYKNLYSEDEGSKLV-QKLP-ELPPSNSDSVSPVTKSVAFSDRIESSPV-   128
9   PTPKRRKLLSPSISENDQLDL-------SKSHEHSSLEDVNNKLM-QSLP-EISASNSNNVSPVTKSVAFSDRIESSPI-   128
10  PTPKRRKLASSSNCENKQSDL-------STTNTNSYSEDAGNRLM-QSLP-ELSASNSDNVSPVTKSVAFSDRLESSPL-   128
11  PSPKRRKTRSLSPSKIAEP-R-------NRHAG-TF------------AD-ENRCGTGANSSPVKKSVAFSDKLESSPT-   121
12  PSPKRRKTASLSPIRNVESTI-------DGPSN-TF------------PI-EARVVTSVQSSPIKKTVAFSDRVESSPT-   124
```

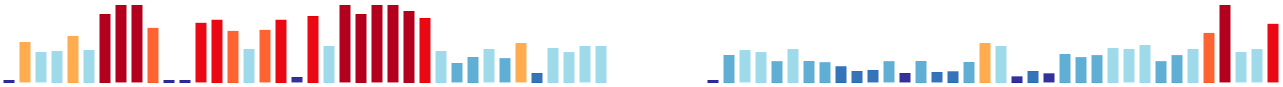

```
   -XXXPXSSPR--PSXXSK-PXKSILRNXXXXXXXXXSD-------XXXXKXXXXXXXLXXXXIXXXXXXXXXXXXXPXXL

1  -YRIPGSSPK--PSPSSK-PGKSILRNRLPSVR-TVSD-------LSYNKLQYTQHKLHNGNIFT-SPYKET-RVNPRAL  193
2  AQTNSAASPR--PSSTSKLRSKPILKLNRVSLQKGDIV--------TE-------Q-----RL--TFSARTTFSLNPMEL  182
3  -STSSNSSPR--HSPTSK-PKKSILRNSSSPEHNKQRM------NDKYGKPIAANV-----HL-SRSENHSKNQYSPDNL  217
4  QSPMPDSSPRRPPSSATKPPGKSILRNPDFITNRTKSD------------------LLSNKDK--LVDSMSPLQLDPGNI  187
5  TQTLPESSPRHPPSSDRKPPSKSILRNSKTPSSPVKYNTIKASSDVTYSKTVS--NNLAASNA--GVATNDFLHADPRKV  210
6  -RITAKSSPT--PSSTSKPPVKSILRNPGATQD-TDEE-------LNHEKSQR-------DSI--VPIDNDSSETNPNNI  189
7  -QRTMNSSPR--QTPASA-PSKSILRNPIEFQK-NVSL-------MADP-AGKPKSKLA-GR-G-GQVNDTSDDIDPTSL  188
8  -HHVPGSSPK--PSPSSR-PGKSILRNKMSSAR-TVSD-------LSYNKLQHPEHRLHNGNILT-SPYKETIVINPRTL  195
9  -HHVPGSSPK--PSPSSK-PGKSILRNRMPPMR-TVSD-------LSYNKLQHSQHQLHNGNIFT-SPYKESTGVDPRTL  195
10 -HHVPGSSPK--PSPSSK-PGKSILRNRMPSVR-TVSD-------LSYNKLQYTQHKLHNGNIFT-SPYKETIGLNPHSL  195
11 -QKSPRSSPR--PTSVSK-PSKSILRNNADTFRKSTGD-------LTAKRSELMLT----SHIGSARLENNTSPTDPHSL  186
12 -QQTPRSSPR--PSSLQK-PVKSILRNT-DALRKCVSD-------ISFKRPPFLNQ-----NIDR-NKTDRTAGIDPHRL  186
```

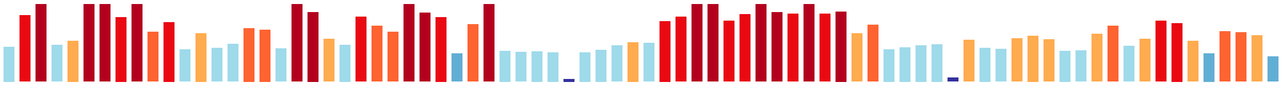

```
   XYWVSGEIHSLXDXXNVXEFKXIIEGGLXILXXXX-XXXVXXRFEXYATFNNIIPXXXX-KXXXEXXXXKXNVLIXNXEX

1  EYWVSGEIHGLVDNESVSEFKEIIEGGLGILRQES-EDYVARRFEVYATFNNIIPILTT-KNVNEVD-QKFNILIVNIES  270
2  AYWRSGEIHNMTDIKNINELKNIFEGGLSLLAVSE-PEHVSKRFEIYATFNAILERQFGDNTPTGMEDKVCSVLCAHMEE  261
3  QFWGNGEIHSLSSNNSVAEFEKLFLGALKILMQSKNPEYGAKKFEVHATINNIMPSFSS-TTSNQFIEKAAHIVKDNLKI  296
4  EYWVSGEIHSLFNNENVKEFEGIIIGGIHILQLENVENSSNRRFEIYATFNNIIPILPTDKDIDPITENKVKVIVNNLPE  267
5  EYWADGEIHSLLYPNNIKEFRDIIEGGISFLNM------GLKKFEIFATFNNIIPIIPNGFNATDVIDKKIGIIVENLDN  284
6  SYWTGGEIHSLKDHNNLAEFKQVIEGGLYVLSSNS-DPHNSRRFEIYATFNNLCRGLSASDHVSETNEKKINILFQKLET  268
7  AFWEFGEVHKMLDSSNFKEFQKIIEGGLEILANNE-RSHVARRFEVYASFNNIIPPNNL-KYHSELIEKKFDILIHNLKK  266
8  NYWVSGEIHGLIDNESISEFKEIIEGGLNILRKES-EEYVAKRFEVYATFNNIIPILTT-KNVNEVD-QKFNVLIVNIEA  272
9  DYWVSGEIHSLIDNESVSEFKDIIEGGLGILRQES-EEYVARRFEVYATFNNIIPILTT-KNVNEVD-QKFNVLIVNVET  272
10 NYWVSGEIHSLIDNESVSEFKEIIEGGLGILRQES-EEYVARKFEVYATFNNIIPILTT-KNVNEVD-QKFNVLIVNIES  272
11 DYWVSGEVHSMLEFNNVHEFKNIVEGGLKILSQTD-AVSITKRFEIYATFNNIMPVISG-KGSSEIAERKINVLIDHMDI  264
12 DFWVGGEVHSMLDPYNVHEFFNIIDGGLRILSEKT-DECVVKRFEIYATFNNIMPPSST-KGLNDINERKVNVLIDQMEF  264
```

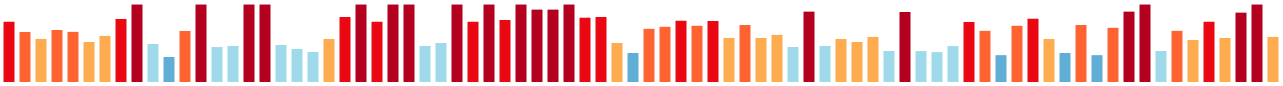

```
   IXXXCXPHLXXEQXXLLXXXEKKNPFXXRLYVQIVRFFXXILSNFKIXKWLXKXPXLXXXLKXIYXXSXEALXNENSNKX

1  IIEICIPHLQIAQDTLLSSSEKKNPFVIRLYVQIVRFFSAIMSNFKIVKWLTKRPDLVNKLKVIYRWTTGALRNENSNKI  350
2  IVNVCYPHLKLEQSSLLENVSVKNPFSTRLYIQIVRFFTHLLSNFKVICALSSKRNIQTSMKQVFESSYEALSSPHCNKQ  341
3  IANISIEHLRDEQNSLLQDKEKKNPFSIRLYIQIVRFFTNLFGSFIINKFLISSKALCERMKQVYELSLEALKNENSNKM  376
4  LIKVATLNLQEEQTELLKSHNKKNPFITRLYVQIVRFFGSILSNFMIIKRISTDPELQKELRIIYKISVEALSNEYVNKM  347
5  LIKILISNLKDEQKELLEGNEKKNPFTSRQYVQIIRLLGIMLSNFKIICWLDKRKPLQVMLKEIYKLSEEALVNENSSKV  364
6  ITDVCLPHLDWEQKKLLGTTTKKDPFSSRIYIQIVRFFTILLSNQILVRRLSKRKTLIAKLSTFYEYSIDTLSNSNSNKV  348
7  IIEVCLPHLRTEQEKLLGAGNKKNPFGSRLYIQIVRLFGSLLAHYRVMRALTKLPSLKEQLGEVLELSKEALVHPNANKV  346
8  IVEICMPHLQTAQEKLLSSNEKKNPFIIRLYVQIVRFFATIMSSFKIVKWLMKRPDLVNKLRVIYSWTTDALKNENSNKI  352
9  IIETCMPHLQMTQDTLLSFSEKKNPFVIRLYVQMVRFFGAIMSNFKIMKWLMKRPDLVNQLKVIYKWTNKALKNENSNKI  352
10 IIEVCIPHLLTAQDTLLSSSEKKNPFVIRLYVQIVRFFGAIMSNFKIVRWLTKRPDLVNKLKAIYRWTTDALKNENSNKI  352
11 VVDTCLSHLEEEQEKLLSTNSKKDPFVSRLYVQIVRFLNSILSNFKIIKWLTNKPQLQLKLKRIYRYSIEALTHNNTNKV  344
12 IVNVCLPHLKEEQQKLLSTGEKKDPFSLRLYVQIVRFFNFILSNFKLLKWLSNKPQLQTNIKSIYEFSIEALTHDNTNKV  344
```

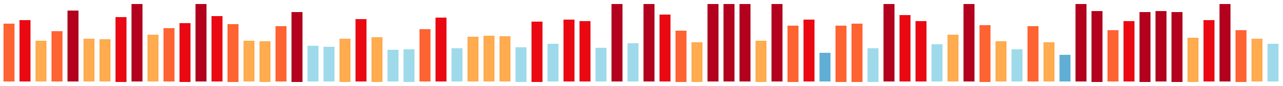

```
   IIXAQXXFLRDEKFGTXFLXXXEIXXIIXXXTXIXEIXSXNLIXEKLXLIRXFLXKYPXLMIXXVXXWLPGEVLPRIXXX

1  IITAQVSFLRDEKFGTFFLSNEEIKPIISTFTEIMEINSHNLIYEKLLLIRGFLSKYPKLMIETVTSWLPGEVLPRIIIG  430
2  MITAQMIFLRQEKFGTSYLTENERNKLVNIITEVNEIDSYNLKYEKLLLIKALFGKYLKQMLQDVPKWLCGEVLARILVE  421
3  MLICQMTFLRDEKLSYISFTEDEGKHMIQIINQIRDIPSTNAICEKLMLIKQFLIKFPGPMISTIPEWLPTQVISKIVFS  456
4  MIISQLTFLRDEKFSRRYLPMSEITKMINVITQVKDMASTNLICEKLYLVKCFLDKFPVLMIDEIKVWLPGEVLPRILLM  427
5  IIISQLTFLKDEKFSTYYLNKNEILRIIDTFTKIKDLKSTNLMNEKLLLMRCYINKFPKLMIPKISDWLPGELLSKIVLD  444
6  MIISHISFLKEEKFGSYYLPSHTIDNLTSIVPNIKEVDSVNLSTEKLMLLRRFLEKYPDMMFNKASLWLPGEVLLRILLD  428
7  IIGAQYTLLAYEKYGAYFLQKEDVMSIIRSVIRTKDIQSTNLICEKLHLMRKFLSKYTSVMLEIIPEWLPSEVLAQILSD  426
8  IIIAQVSFLRDEKFGTFFLSDEEIKPIISTFTEIIEINSHNLIYEKLLLIRGFLSKYPKLMIETVTSWLPGEVLPRIIIG  432
9  IITAQVSFLRDEKFGTFFLSDEEIKPIIDTFTEIIEINSNNLICEKLLLIRGFLSKYPKLMIETVTSWLPGEVLPRIIIG  432
10 IITAQVSFLRDEKFGTFFLSNEEIKPIISTFTKIMEINSHNLIYEKLLLIRGFLSKYPKLMIETVTSWLPGEVLPRIIIG  432
11 IVAAQVAFLGDVKFGTFFLNDEELTSIIHTIPTIKEIQSTNLICEKLILIKNLLAKYPRLMIENVSIWLCGEVLPRILID  424
12 MVAAQVALLGDSKFGSFFLNDDEITSLIRIIPSIKEIQSTNLINEKLILIRSLIAKYPRLMIDQVSIWFCGEVLPRILIE  424
```

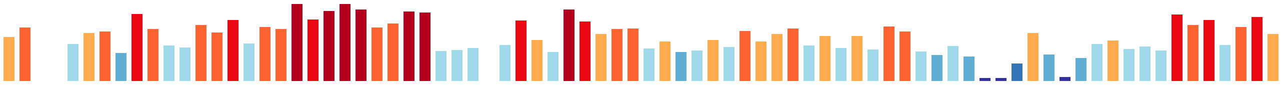

```
   XE--XYSXKIXXTXIXVLLDLLKKCLDXXX-XHEXIYQCXXXXXXXXXXPXKXLXKLXXXX--XDX-XXXXXXTLGXLLR

1  DE--IYSMKILITSIVVLLELLKKCLDFVD-EHERIYQCIMLSPVCETIPEKFLSKLPLNS--YDS-ANLDKVTIGHLLT  504
2  TQ--TYANKLAVVALEILLDLLKQLLGGKDSVTASIYESMEEKPAKDVLSPSMKMMLQSVL--PDT-YDFETITIGSMLR  496
3  ED--VYSTKTGFTAISVLLELLKRCIDHSI-GHEYVFQCLHIDSTALYFGDSKDKKLQWSNNTLEQTYDINETHLAAILR  533
4  EEENNHSAKIIVTAISIILDLLKKCLDVEK-GHKDIYQCVEIDKVKDVIPRKFLSRIELSL---KEIENVGEKTLGTLIR  503
5  ED--MYAIKITHTSIAVLLDLLKKCLDVEE-GHQEIFKYVEVKTLNESVPEGTLEKFNSTAVLNDLEIDINSMTLGQLLR  521
6  DG--SNGWKCIIAAISVLLDFLRLCLDHDN-TQINIYQSIEVDKLNDVVPQKFLPKMQQYL--KDK-SLLESMTLGQVIR  502
7  EE--VHSLSILQAGIAVVLDLLKKSLDSSA-IHQTINRCIRNTLAKDVMPEGAQNRSVHG-----HSLEWGSTTLEEMLQ  498
8  DE--VYSMKILITSIVVLLELLKKCLDFVE-EHERIYQCIMVSPITETIPEKFLSKLPVSS--HDT-ASLGKVTIGHLLT  506
9  DE--VYSMKILITSIVVLLELLKKCLDFVE-EHERIYQCVMVSPVTETINKKFLSRLPLNS--NDA-GDLNKITIGHLLT  506
10 DE--VYSMKILITSIVVLLELLKKCLDFVD-EHERIYQCVVLSPITETIPEKFLSKLPLNS--SET-ADLSKVTIGHLLT  506
11 SE--INCSKIVLTAVSTILDLLKKCLDVSK-GHEDIYNSVQICTVREVVPAKLLPKLLPSQ--NDP-ESIISQTLGQLLR  498
12 NE--LNSTKIVATAVSTTLDLLKKCLDFSK-GHEEIYRCVEVLFVKDVVPKKLLTKLTPSQ--DDNLDSIAQQTLGQLLR  499
```

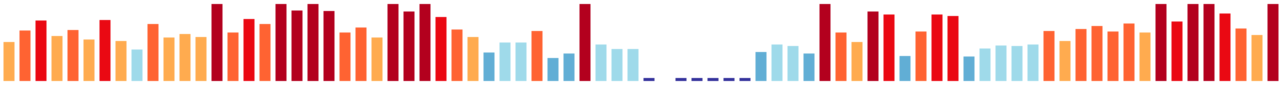

```
   XQIKYXIXXKXXYKXAMDLWLXMXGLLYNXXXXXXXLXXX-------XXXXWXXLNXLCFXXXXXKXXLXSXKVWRIXTY

1  QQIKNYIVVKNDNKIAMDLWLSMTGLLYDSGKRVYDLTSE-------SNKVWFDLNNLCFINNHPKTRLMSIKVWRIITY  577
2  DHLRNQITEQKAYKIGIDLWLALVGILFNSPRNITKLLPT-----DDRGDSWLSLNELAFKTNDEKAMLLALKSWRVVIY  571
3  DHIDYLVNSKREYKLAMDMWLSFMGLLFNSKGRLAFLASK-------EGEKWVDINLNCLSVNDPKCRILSLKVWRILIY  606
4  NQIKYLILVKKEYKLAMDLWIGMVGLLFNNANSLEMLTKENAETDANRSDTWIDLNKLCYNKGTGKVKYFSSKVWRILFY  583
5  NKIKYLVLVQKDYKVGIDLWLAMVGLLFNNKKNIIHLLDKD-KTHDKSSADWLNLVQLCFKNSDGKTKLLSLKVWRILTY  600
6  DHINYFLLANKHYKIAADLWLAMVGLLYRKADKIEELSAGT------YENNWLSLNSSCFEIPEEHIKLMSLKVWRILFY  576
7  KEITFLIISKKEIKFAMDLWLAMMGLLYNTTEALLELSDT-------DERGWLRLNHICFLSGNSSAKFIALKAWRIITY  571
8  QQVKYYIVHKHDNKIAMDLWLSITGLLYDNEKRVYDLTLE-------SNEVWFGLNNLCFINDHPKTRLMSIKVWRIITY  579
9  EQIKYYIVKKNDNKLAMDLWLSMTGLLYDNGKTVYDLISE-------SSKTWFDLNQLCFMNEQPKTRLMSIKVWRIITY  579
10 KQIKNYIVVKNDNKLAMDLWLSMTGLLYDNGKRVYDLTSD-------SNKGWFSLNNLCFINNQPKTRLMSIKVWRIITY  579
11 KQLKYLILVKQEYKLAMDLWLAMTGLLYNSTDRLQQLTSA-------TDREWLDLNFLCFQADDPQAKLLAIKVWRILTY  571
12 KQIRYLIVAKQEYKLAMDLWLALTGLLYNNIDRLDYLSTH-------IDNEWLSLNLLCFHASSAQAKLLSIKVWRVLTY  572
```

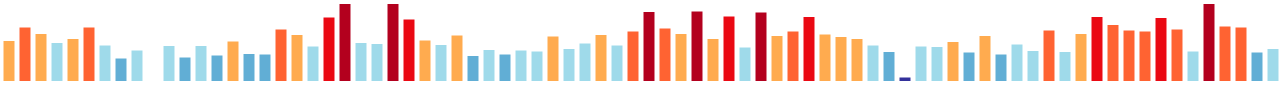

```
   CIC××I×××-×××××××LL×LL××PF××××××××D×××REG×IY×L×G×VYTAF××-××N×S×××F×LFWD×L××PIY××

1  CICTKISQK-NQEGNKSLLSLLRTPFQMTLPYVNDPSAREGIIYHLLGVVYTAFTS-NKNLSTDMFELFWDHLITPIYED  655
2  LVSVNITSLESDEVQLQLLQLLLTPFDQLPNVLEKDCVREGLLYCLNGVVFCSVSMINSLGTQNAISLFWDNLLQPLFHK  651
3  CVTNNIDFL-GEKEAKKISQVLLNPVNVTVNKLEDNSTFMGVKYLLNDFFFSTCGI-TNKCKPTTNIVLWGNFIFPMLAT  684
4  CVSKHLDEF-NETSKPMVLDLLKTPFDMSRPYINDSDVSKALLYCLRGLVYANLCGP-AQSSKESLSLFISNLLKPILIG  661
5  CICTHVTEL-DSAYQIPLLNILKIPLEISKTEHGNDNVRDGIIYYVTGIIYTAFLLLQKNLSYETFQFFFDNLVKPIYEN  679
6  GVCSEISGK-FDEKDLLMLNLLRRPFELSEGQHTMPAVRDGLLYCLNGLTFTIVSTVNVNNDAKRFGFFWDNLLKPLFIS  655
7  YSWAHVQGK-SLHHDHALIRLLKMPFEFSSTSHLEPDVCEGLIYYLTGMVFVTCGN-SKAQEQKTFRFFWENLISPIYHD  649
8  CICTKISLK-NQEGNKRLFSLLQTPFQMTLSHVNDPSAREGIIYHLLGVVYTAFTS-NRNLSTDIFEHFWDHLIAPIYEN  657
9  CICTKISLK-NEERNESLLSLLQTPFQMTLSHVNDPSAREGIIYHFLGIVYTAFTS-NKNLSMDMFNLFWDYLIAPIYEK  657
10 CICTKISLK-NQEGNKMLLSLLQTPFQMTLPYVNDPSTREGIIYHLLGVVYTAFTS-NKNLSADMFELFWDHLIAPIYED  657
11 SICSHLETF-S-TENLRIIKLLQRPFTFSREDQYDFNIMEGLLFHLTGIIYTAFSA-TNNLSTAKFDLFWEHLVRPIYVE  648
12 SICTHLEHE-T-VKNIRIVNLLQRPFEFTQQEQYEFNVMEGLIFHLTGIIYTAFSG-HSTISGEKFSLLWEHVVKPVYMR  649
```

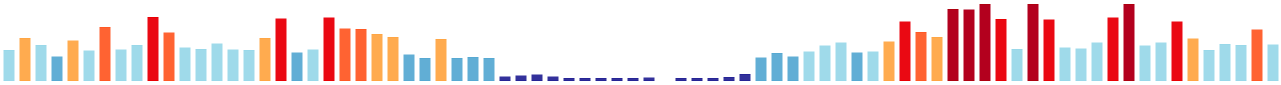

```
   ××F×××SI×L×××××××L××L×GGK××××××----------------×××××K××HP××VIAS×GV×××DI××LP×××I×

1  YVFKYDSIHLQNVLFTVLHLLIGGKNADVAL----------------ERKYKKHIHPMSVIASEGVKLKDISSLPPQIIK  719
2  WNSAASSNELQTRTMDIFIKILRGKFPKTSTQRSNS---------PNTKQQQHNFRPVRVIAARGIVISDIPNLSNNFIH  722
3  LLNYSANEDVKFELDTYIGKLLTKAIQN-----------------QKNVDKTKSLHPIKVIASEGVSVDDISALQLSQLP  747
4  IILVNQH--LHHEVVSVLLRLLGKPLPETRSPEK------G-------NRNGLRPAPIKVIASTGIATNDISKMTNHLIK  726
5  FIFNSPSFHVQTRLLGILLQLLGAPLFNLVSTSKHITSNAGKSKNIRNDKKVSPNNPVKVIASVGISRDHIVPLPTHLSH  759
6  FLKTSPNNQLRARVIHLLALMMGGHGFDVVKRKRRPGDQEP-YTTKKQKVIRAYADPVRVVAATGISLDDITSLHPNVLN  734
7  FLFTSDSIKLKSAAIKLLAVLIDGKSETGTS-------------------KTKSSFPLKVIASVGVTMKEIEPVPADTVD  710
8  YVFLYDSIQLQNILFTILHLLIGGKNNDFAT----------------ERRNKKHIHPMSVIASEGVKVKDISSIPSQIIK  721
9  FVFNYNSIQLQNVPLTVLHLLIGGKNSDVVV----------------ERKNKKHIHSMSVIASEGVKIKDISSLPPSIIK  721
10 YVFKYDSIHLQNVLLTVLHLLIGGKNADVAL----------------ERKNKKHIHSMSVIASEGVKLKDISSLPSQIIK  721
11 RILISQSIQLKARANALLLRLLGGKITEPQH----------------KQVCKKIVHSIKIIASAGVTARDIQPLPLNVIK  712
12 KIFPSTSIQLRNRAIVLLLRLLGSKVMEQQANQQHQQEKQQ-P-HTHRNITKNGAHPIRVIASAGVSARDIQPLSANVIG  727
```

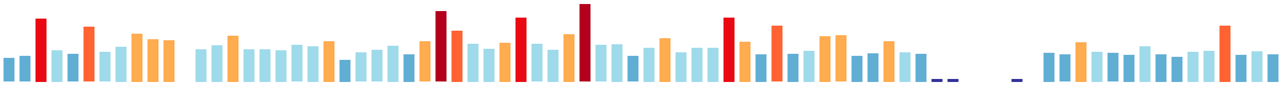

```
   ××Y××I××××F-×××××××SN××××××LIL××L××LP××××××××LE×F×××××××××-------××××××××××F×××

1  REYDKIMKVVF-QAVEVAISNVNLAHDLILTSLKHLPEDRKDQTHLESFSSLILKVTQ-------NNKDTPIFRDFFGAV  791
2  IWYMQMAKSAS-DLLRTRSFEDEHAKSLFLSLIDTVPESSLNDEVFDYFLLVLKHYLEP------TYDYTFNQLSVLTLF  795
3  YLLETLGSVILLYVLNSKSQNHSQTRMLILEYLEQIPNNQITLSTLCRCISMFYEYYN-------SIDFIANSPRSLISD  820
4  QLYYPLSDLIF-SILRKHILKPQPSSELLISLLNNTPLELKTVSYLEEVNQVMIDILKT-------SDDHTEISLLFQTY  798
5  PGYTLISRLFL-NSLPINAKFLTVKTEIILDLIAQIPDELKTEEVFQEYRNIVVSLTKLYKTSYTTKNKNDVLFECFSRC  838
6  ENYLSIMGLCM-DNTGAHPLS-YLTDKLLYLLLSKLPEHLRNTEVLEAFSDVLSFVVNRS---V-RISSKANTVRIFSTH  808
7  DAYELVLTLVS-NALSIDSSEWATRFELFDSMLRLLPAEFIDDTHLTIFLDLVSRLFS-------QASLPKDPSTVLCPL  782
8  YAYDKIMKVVF-KAVEVAMSNVNLAHDLILTSLKQLPEDKKDQNHLEAFRTLILKVAA-------NNKDKPIFRDFFVTV  793
9  SAYDKIMKVVF-QGVKTAMSNVNLAHDLILTSLKHLPEDRKDQSHLEAFSSLILEVAE-------NNKDKPIFREFFGAV  793
10 SAYDKIMKVVF-QAVEAAISNVNLAHDLILTSLKHLPEDKKDQIHLEAFSSLILKVAQ-------NNKDKPIFRDFYGAV  793
11 SSYEAIKDLVF-QAIESDSSNLCQNFDLITALLKVLPNKFIDKEHLREFTNAIFGVAI-------QRKEDQNLADVFIQI  784
12 TNFETIKDLIF-EAIRSNSSNLSQNSELITALLKQMPSKFVNENSLEDFAELIREISI-------QRKDDKNVGSHFVQL  799
```

```
   XXXXXXXFXXLXLXXXXXXXXXXXXXXFXXXXXXXXXXXXLXLLKXXIXX-XRXXXSEFXIIEXFLX-XXDKXXXXYXXNWV

1  TSSFVYTFLDLFLRKNDSSLVNFNIQISKVGISQGNMTLDLLKDVIRK-ARNETSEFLIIEKFLE-LDDKKTEVYAQNWV  869
2  ATQLVTKFEKLLLEPSNSGLSSVIALYDSKTFINYDFPLKLMISMLDS-NDCDASLFAIFSKFMD-MKNKNYPDYVTNWL  873
3  MSRVCIKFKDIIFDDNYNVFRNFILSFLMLFQEEEHHVDNLLQELIQNALQANISELKLIEGILS-LKKTDCDLFAINWV  899
4  THVVVTIFPNLLLNKETNYLEDYLKLFEPFWGDVKEMLLDVLKDFLRE-KKLAGWEFYMMGIFLK-TKDTLSHLYVENWI  876
5  TCLLGKTFPNIILNPKRDIFSKYLASFDNLFTDNADMLIKLLKDIIKY-TRPQISELTVIELFLTNTNDKNSRDYIQNWV  917
6  SSKLAITFKDILFDKNSKAFTNYIAKFNMLSDREGTCLLTFLKQLLNN-SKGIIPELFVVENFLL-IGNKEVDSYLSNWV  886
7  VLSLTGPFANFVFESSK-NFQKLLKGVDHFYGQDSEIRLTLMKEFSKK-LRRQISEFTIIERFLE-VGDLSCQRYASNWI  859
8  SSSFVYAFLELFLKKHDSSLVEFSIQLSKVGIPQGNMTLDLLKDVIKK-ARNETSEFVIIEKFLE-LNDKKTEVYAQNWV  871
9  GSSFIHTFLDLFLRKNDSSLINFNIRFSQIGSPQENTTLNLLKDVIRK-SRNETSEFVIMEKFLE-LKDKKTEIYAQNWV  871
10 IPSFVYAFLDLFLRKHDSSLINFNIQFSKVGILQGNMTLDLLKEVIRK-ARNETSEFLIIEKFLE-LEDKKTEVYAQNWV  871
11 TCCLATQFATLFSDERK-IFEEYLGKFSFILLANGETKIKLLKELIVG-LRGKMSEFFIIEAFLQ-QGDLSTKEYVSNWI  861
12 TCCLMSQFTPLLFEDRK-NFEAYLRKFEFIFSVNPDTKVKLLKELIMG-LRGKVPEFFLIESFLE-PQNGAIRKYVSNWI  876
```

```
   GSTLLXPXXXXXEFXSLANIVXXXPXXXXIENFLDLCXKXXXXVNXXXLLXXXXWXXXXXXXFIKXYXSKNXNXLXXXLX

1  GSTLLPPNISFREFQSLANIVNKVPNENSIENFLDLCLKLSFPVNLFTLLHVSMWSNNNFIYFIQSYVSKNENKLNVDLI  949
2  SVAVLPSNMNRHDMKFLMNIISKNPLVSIIENLLTVIADIDREFYGMICTSALTWDDHALLYFIKGVFTSNAPNSCNYL-  952
3  GKKLLSPSVQEGDFILYSRIVAKISEKQTIVNFLDLADKLGFEVNPTEILRISFWKPSGLVIFLEEYIAKHHVDFADIVS  979
4  GSTLLSPTMSYTSFLDLANIINQITTDQVMENFLDLCSKMSLSINYPQLLNLKQYNDKNLSFFIKTYLSKNSNALDNNMS  956
5  GSTLLSPMLTYPEFQSLTNIVNMIPSSQIVENYLDLCVKTSLAQNPCTTLNIRNWKDKQLIFFIKTYISKCSNRLDDDIL  997
6  GSTLFPPSLSKDEIIALAHCVSTIHTEKAIENFIDYSKKIE-QLDVVTLLHLGAWDESDFCFFVKAYLAKRKNVSVTHLI  965
7  GSVLLPNTLTTSELHSFASIVKLLLSPQAIGALLNFCLNSKDKPDICELLNIDDWGDEKVAILVKAVVSLSSNKLVPSLA  939
8  GSTLLPPNISFKEFQSLANIVDKVPNENSIENFLDLCLKLSFPVNFFTLLHVSKWSNNNFIYFIQRFFSKNDEKLNADLI  951
9  GSTLLAPNISFKEFQSLAGIVDQVPNEHSIENFLDLCLKLTFPVNLFALIHVSKWSNDNFIYFIQSYVSKNNNKLDTDLV  951
10 GSTLLPPNISFREFQSLANIVDKVPNENSIENFLDLCLKLSFPVNLFTLLHVSKWSNNNFIYFIQSYVSKNENRLNADLI  951
11 GSMLLSPTMPHEHFSTLVNIVTLIPTTEVLENFLNLSSKVNYDVDLFSLLNIQNWEFQGIIIFIKNSILKNHNTIKPELN  941
12 GSMLLSPTIPYEHFCSLVRIVNAVPTTEVLQNFLNLTSKVPFEVDLFHLLNLKSWDDKELTLFLKNSIIKNLKPNRPELY  956
```

```
   XXLXXXLPXNX----XLFXXLLPXLXXXKXXXIXXXXIXXNPXXLNXXXXXXXX----XLXXXLPXXXXSYFXXNXXXF-

1  TLLKTSLPGNP----ELFSGLLPFLRRNKFMDILEYCIHSNPNLLNSIPDLNSD----LLLKLLPRSRASYFAANIKLF-  1020
2  ---SGCLPTRISKSLFLFKAILNSPELFKENGIVLRSVLKNPSLIQNLNVAEVT----KLRDLSDHDAPTDFMKYIN---  1022
3  SIIDADRLNNV----ELFSYVLKHLPENQKSAFIKQHYTKCPLIWNQGVLIDDY----DLIDITLDCEICTFIAHHFSFL  1051
4  QFLIEILSENRTS--SVPRILLPLLIDIKRYDIIKTSIEQRPLILSIVSDPILL----HLKDILPTSKMSYLIINIGLF-  1029
5  VLLHDVLPDNI----DVFDSMVPLLIQINRTDLIRHCIEKKPSYLNTPAIMNYH----DLSEILPDDRKCYLLNNLKDF-  1068
6  ALLQKLLPSRP----ETFTELLPTLLEYGHKELVSDCVKKNPILIESLLPKYPD----NLSDFFSADEILAFVPDMKNF-  1036
7  VKLQKMLPSRV----SLFNELFTTLQTLKYEELIRHVIIANPEAANGIILLNGTQAPAVLPKGLPRDTLSYFVSQLPNF-  1014
8  TLLKTSLPGNP----ELFFGLVPFLRLNKFIDILKYCICSNPALLNSITDLNSD----FLMQLLPLSRASYFAANIQLF-  1022
9  TLLKTSLPGNT----ELFVGLLPFLKEKKFMSILEYCIQSNPTLLSYISDLNSD----LLLELLPRSRTSYFVTNIQSF-  1022
10 TLLKTSLPGNP----ELFFGLLPFLRNNNFIDILEYCICSNPTLLNSIPDLDSD----FLIKLLPRSRASYFAANMKLF-  1022
11 LFLQSVLPNST----PIFSEMLPFLCGAKHYNIVRYTLTKSPKFLNLTTSVDKK----SFDEILPLEILPFFLQSISDY-  1012
12 SFLENILRENE----QLFSTLLPLLCQLGHYEVLRLVLTHRPTFLSHTLGVDEE----SFDQILPLEKLSFFLKNLLNY-  1027
```

```
   XXXXQLXXIXWXLXXXXXXXLXXNXX---------XXXXXLXXX-XXXXXXX-XXXXXXIXXLLXXXXXXXXXXLLSXLXX

1  KCSEQLTLVRWLLKGQQLEQLNQNFS---------EIENVLQNA-SD---SE-LEKSEIIRELLHLAMANPIEPLFSGLLN  1087
2  ----------VVYKKQIAGTMNNKLSQSGTSTDKSNSECFNEPQLDNQGQTGEDTSVVVLEKLHECCSNNKGDALTEILS  1092
3  STSSKAKLLIIFIEKDMYDLLVQFRY--------NLIQEFLPTLTIYKKD---DIITTVQQHLINYCFAKNSFVLLDCIEP  1121
4  PQLIQYGIIQWVLEKDETALLLSNYA---------TIETILFVD-KPTDTSVNEERKKLIQHILHYLFENRRWVLLSNLID  1100
5  SAIVQYKIIMWVAKTDEFTLFKKNYV---------KIESCLFLD-EPCSENILD-KKAFIKELFTIFYEKQAWQLVAILVV  1138
6  SSETQVILLKEAWEKSGLSTVVKNFD---------SVASCLFEQ-ALDG-SISNVRKILTDSILSSIIKDERWDLLNNFFV  1106
7  DENAQLGMIKWILDMNEADLLFADRG---------EWKDCVFLR-IGKDGEFSNEFKILIADAISQLYEKQSWKNLSKCIL  1085
8  NNSEQLTIIRWLLKGQLLEEINQNFS---------EFKSSLQNA-SG---HE-LERSEIIRELLLLTAKNPIEPLLGGLLE  1089
9  NHAEQLTIICWLLEGEELDQINQNFP---------EVKNILLHT-PA---DE-LEKSKIIGELFRLATKNPIEPLFSELLN  1089
10 KSSEQLTLIRWLLKGQQLDQLNQNFS---------EIESVLQNA-PD---SE-LEKSEIIGELLCLTTRNPIEPLFSGLLY  1089
11 QDTIQLRILQWCIEHNQINMLFGHYE---------TLEHYLFKK-EYGTEES-SRRNQLLLDLLSSLLVGKKWLYMSKIIQ  1082
12 ETFFQLRTIRWALKHNQINILFENSS---------VLQKCLFQQ-CTSSEDT-MKKEKLLLDLLSVTVLGEKWHYLSLLVI  1097
```

```
   XCXXN-XXXXXXXXFXX---XXXXXELXXXXPXXLXXXXXXXXXXNXXLXX-XXXXXXXX-XXXDXXXELXXXLIXXKXX

1  FCIKN-NMADHLDEFCG---NMTSEVLFKISPELLLKLLTYKEKPNGKLLA-AVIEKIEN-GDDDYILELLEKIIIQKEI  1161
2  NLLSQ-DQTTIISNFFCSYAAQNIEVIKLISPKLLRHLCKFYPDHLHALILPAINYLFGPVQKISTLTSLAIILIEKKDI  1171
3  STLTS-RTLEDVIKLLE---KSYEDNITFVPAKYIIFIDNLPMDTKINTTE-ILRVYLVT-GDVTQCLFFLTSLINDKKM  1195
4  SCLSN-SQTDYIISLFS---GGNATELQNILPQTLANMINRCGSLNPSLID-TLKQNYSL-QKADYIITLISELVECGKT  1174
5  SCINH-GYESNATEFLA---NIDSNTLNSFPPKAIATIVNRCGSMNPKLLL-ILHQFFSN-KEKFFLLELVSELISLKKI  1212
6  TAYLNGRDTGLLDNFFT---KVDSESLEHFQPNFLVAICDRLGY-SERRKN-VLKKCFFV-KEKEYVISLIEALIKSKKF  1180
7  YCMTC-GQVPCVEQVYV---RMRDKRRENNSPK--SPKADECKSLNSILSE-SIKEIYQG-DAIDTAFEVTTSLIELKKV  1157
8  ICVEN-SIIDCLNEFCE---AITDELLLKINAELMIRLLTYRDNPNEALLT-AVTERIRN-ADSDYTFEFIGKIVARKEI  1163
9  FCIDN-SMTNNLDEFCE---TINDEILSKINPELLMRLLTYRDKPNVTLLT-VLAGNIGN-ADSDYKFELLERIIARKEL  1163
10 FCLKN-SMTDHLDEFCG---KVTDEVLLKINPELLLKLLTYNDKPNGKLLT-AVTEKIGN-ADNDYILELLEKIIAQKEI  1163
11 SCMKN-GQASPVTKTFS---KNDPESLFWLTPVTLASMINKCGLLNPSLIE-VLKKCFSR-KPADFNIELTKELINRNES  1156
12 WCMEN-NEIGHLTTCFS---KENLTPLIHLSPQALALMINKCGSLNPTLIE-AVKRSFFT-NDTQFNIGLIEELIVHNKF  1171
```

```
   XXLXXXXXXXXXFFXXXXXXXXXXXXXXXXXXXXXLXXXLXX--XXXXXXXXFXXXLXSXLPXXXXXXXXXXXXXXLXXXX

1  QILEKLKEPLLVFFLNPVSS-NMQKHKKSTNMLRELVLLYLTKPLSRSAAKKFFSMLISILPPNPNYQTIDMVNLLIDLI  1240
2  NGLDRIIDPFSIFLFKGPKKLSSDIKHKIAGLFLKMFFLLKE--ERGVHLSIVLHRIVRSLPRRRPEYLKNVINTISTQK  1249
3  VPLTRCRNEVFDFFIALDSSNDLSIKAKAVTTFESLINVIFL--LRKKVDIGFLSQYIARLPERPSRYLVDLSEKLILNK  1273
4  QALILSKQDMLLFIIDSENALRPVDQRRILELMPRFIQSLQN--KSEKTFIDFLKMLFNLLPEHSNQYLLNLVQLFKESL  1252
5  DALLSVSSELINFFIDDQHIFTEDEQGKSIGILEKFLAQLQK--YNERTSFNFTKDLMSSLPKSKTKYLSKLASCVGSAQ  1290
6  DVLNDLRSEFITFLLVHSKDPSLTFEKQYVDGINNFILLLKE--QSEDVVVDFMVYLISPMQKQMSINDKTVLKYMISDE  1258
7  QNFSSCREEFLNFFIIRASNFVSVEQAIALELFERFIAVLME--CFDEIILSFVESILSSILKVLTDHSLQVFGLLNHHL  1235
8  LLLEKLKEPLLVFLLSPVRS-SEQKYEESANMLRELVLLYLTKPLTRSAAKKFFSMLVSILPSNPNHQTIDMVYLLIDLI  1242
9  LVLEKLKESFFIFLLDPVSRLNKENHDNGTKIFCELILLYLTKPLSRGAAKKFFSMLVSTLPERPDRQSIDLVHLLIDII  1243
10 QLLEKLKEPLLVFFLNPVRS-NKQKHEKSANMLRELVLLYLTKPLSRSAAKKFFSMLISILPPNPNRQTIDMVALLIDLI  1242
11 QVFFLCGKEVVAFFTVRSKMFTLSEQDAISALFEMLMTSLMC--QKDSLILDFVQELYANIPQECDSHLLKILIIAVKKL  1234
12 QLFILCGKEIVTFFLDNSSSLSGVEQDGVLSIFEKLLVSLMG--QKKSLVLDFVEEISEKLPEAPDAYLLKIITILVEQL  1249
```

```
    XXXXXXXKXXXXXXXXXXXXXXWXXXXXXXX---XXXXXXXXXXXXXX----XXXXXPXXXXX--X--XXXXXXXXXXSX

1   KSHNRKFKDKRTYNATLKTIGKWIQESGVVH---QGDSSKEIEAIPDT----KSMYIPCEGSE--N--KLSNLQRKVDSQ  1309
2   NLE----NKIPDINDIQLALTKWIKLEDYSSYLLKYRRYAYLDQITNQVGITQTNRQPHSN-----------GIVGTNIA  1314
3   SFNRSKFRNSSSFNDLHTHVRTWCNQLDNTSGS-----PRETTPIS----------AAHIN----D--------------  1320
4   PSSFKNSKSRREMGRLNKNFQRWKDETKSIEPTSG------EQAISSE----STMSTPE-GFDKST---KSQNSSSNDNA  1318
5   NST--------VVGVTQEELKLFSQKLEDCD----------NQEVNKT----LAIVQPKAQFVVSRDIRDVYTNKIQDSD  1348
6   YIKNVSIKRIPKFDDLVDHLIALE----------KQDSSNLTMTSFSD----SSLSTPEGLQ-------ISNNKNSEESA  1317
7   TLKGHLFHKLENYKTLLEKSGESKLVKENVEAQCSNGGPTLLEPIH-T----GTTSNASDMGQDKN--QSDDINKAAVSY  1308
8   KSHNRKFKDKRTYNATLKRIGKWIQESEVDQ---QEDSRKEIDTTYSA----NSINIPDKESE--K--NVLNSQRGSNSE  1311
9   KSHNRKFKDKRTYNATLKTIEKWIHDSEVVL---QRDPDKEAGEKLND----SSIYTPQGLEVEKG--VPENLQKTPNIE  1314
10  KSHNRKFKDKRTYNATLKTIGKWIQESDVVH---QGDSSKEIDTTPNA----NSTYVPHGESE--K--EVSNLHRKADFE  1311
11  QAKGKHLEKYDRYHHISGELFSILSNLTITS---QNVGLK------------NGIAQETPQ----N--NESN-FTEKNSK  1292
12  QVNGKKLKKENRFSQALQRLVELHYN-----------GIK------------NQSYEDHHE-------NQER-IRETQSS  1298

    XIQVPATQGXXXXXXX---XXXX-----------XXXXXXXXXXXXKXXXXXXXXXXXXXXX--XXXXXXXXXX---XXX

1   DIQVPATQGMKEPPSS---IQIS-----------SQISAKDSDSISLKNTAIMNSSQQESHA--NRSRSIDDETLEEVDN  1373
2   QIEVPEMQSLSKTKDNAAIEEGSAQQGVET-----------------------------------LI---TG-----TNS  1351
3   HINKNALSLEKDKDNL---TNN--------------------TEQQKNIPNSKS--------------------------  1351
4   EIQVPATQLKQDNDVT---TGTTTAKSAYSYQFKSSTEDESTDVLSQKSTGIIKSPEE-------LIQPTKNTH------  1382
5   EIQVPATQKDEEKEII---AIGSQSQSNNDLNNSSIEEENKNNTIVSKNLAIVTSIETPSGTQSQEIRQLTGNH------  1419
6   EIQVMGTQDVNKISGI---SNMR-----------AIH------------SPQVNFSDSRHST--KIIASSPNVVNGQDRN  1369
7   IIEVPATQGISKSAES---TEGR---------------------------------------------------------  1328
8   DIQVPATQGIKDLPSS---LQLS-----------SQNDIKSSHTIKLKKFPITNLAQPEILA--EISRSSTNEFLGEADG  1375
9   EIQVPATQGVKDLPSS---FQTP-----------SQADVEDFYVRKVKNIPIMKLTQREIPA--KLGRSFTDETLEEVDN  1378
10  DIQVPATQGIKDPPSS---YQLS-----------SQLDAKNSGTIKLKNIPIMNLAQQETLA--KRSRSFADETLEEVDN  1375
11  ISMNPQRRGL----------------------------------------------------------------------  1302
12  SENVPTPKGFNDPRIS---ESP---------------------------------------------------------S  1318

    XXXXX-XXXXXXXXQXXXX-XXXXXXXXXXXXXXXXX----XXLXEXXXXXXX---------XXXXXXXXXXXXXXXXXX

1   ESIRE-IDQQMKSTQLDKN-VANHSNICSTKSDEVDV----TELHES-IDTQS---------SEVNAYQPIEVLTSELKA  1437
2   IGMQE------NFSQEPLNDSI-DRTVHSHEGNGTQATYLANALSRIDNEQNHS--------TKQHNASTSNSLTSTLSE  1416
3   --------------QEISK-FS---------SNIADA----NDLPSNTRESNC-----------DG-SQSTEPAGGKMIK  1391
4   --------------------YQ-------EEKAPLVS----QELPSYRSSQDPRSTQEGKITNGDGTAIPDSSIIGTLEE  1431
5   --------------------LE-------NESK--EV----SDLPEVYS---PSDT--------VNKRIDDSSFTQTLDE  1455
6   LAMKKENKVPTISGTEDKKIFVNSSSDIISCSDDAHH----LNLFEGDKNANR---------LDDSSEEQMKNF------  1430
7   -------D-----SQRDMN-IVTSTIIHD----TVE----------------S---------TEISSFVKMDVASQK---  1363
8   ESTGE-GDQQANTTQLDDK-TINDKSIYSTNNGEMQV----TELQEYS-GVHP---------SKANSDQQTEVVIDKLEG  1439
9   ESMGE-IDQQAKSTQLNNE-IDNHKDIQYNKSDDAEV----SELHEDDATANS---------SE----------------  1427
10  ESARE-IHQQMKSTQLDEN-TVNRTSIHFTKSDETEV----SELHEDDNDTHS---------SGANAYKQTEVFTGELRE  1440
11  -ESHDRLEIQVYATQTKTS-QIEE---------PVAV----EEEGKNNEGRRF---------STDHSVAEKKTTTAE---  1355
12  EKIQEINEVQVYATQEKHK-VQNE---------PVKL----SHQTEGQVSLVA---------SEP--SSDK-----K---  1365
```

```
    XXX--XX-----------XXXXXXXXXXX-XXXXXXXXEXXXXXXXXXX-------------XXXXX-XXXXXXX----X
1   VTNRSIK-----------TNPDHNVVNSD-NPLKRPSKETPTSENKRSK-------------GHET---MVDVLVSEEQA   1489
2   VTS------------------YGGLIQSDTQEVRT----------TLNK---------------ESS-IDDDIVD-----   1447
3   ID--------------------KGQISAD---------------------------------------MKNNKIA-----   1407
4   VDKQIAK-------EPVTQSTGCGQLENH----TNIVGESSDLPEVLEDEN-------------AGSIIKDDLLSTENRN   1487
5   VEDPENVNMIQAKSTQIGPTTGDTQINNI----SDTNAEAPDLEDKVDEEK-------------QE--------------   1504
6   --------------------------------VERQEGEKENVKEKFRQGSASKKSERQRDIAHAGSKTN-DRLRPQNDS   1477
7   -----------------------------------A-------SEEVAR-------------EHESS-------------   1375
8   VKKELVK-----------TSLDQNLVDND-DLVQCFPGESPVLEDKRPK-------------DHEKDKKMGDVSSFEEEG   1494
9   --------------------------------------------------------------------------------   1427
10  VTNKFFK-----------PNPDHNSVNSD-NFLQRSAEETPISEDKTSK-------------GHEE---AANVLPPEEEA   1492
11  -----TR-----------ESPQEIIVQSNTNPNK----EAPE--------------------DAASAKTM---------G   1386
12  -----NR-----------PILEEAVI----DPR--------------------------------SG--R---------G   1382

    ----------X--------------XXXXXXXXXX----------------XXXXXXXXXX---XXXXXXXXX---XXXX
1   ----------VSPSSDVICTNIKSIANEESSLALR----------------NSIKVETNCNENSLNVTLDLDQQTITKED   1543
2   ----------L--------------SQKSSPVE---------------------AVGTTCV-------------------   1463
3   ---------------------------------PR--------------QISSIASEPDTL---VNSNIDNKSMIEKGKN   1437
4   DTEIGNELVVAEPTPTVVEAKIATVSEQVIPMTPSKTNPQTDAETPFQTPRSSLGVEPHASEKDAAKTIDSDD----KTN   1563
5   ----G-------------SSNLYTVGNVVLPFACSPEQRGVDN--IQQGEISTLSTSRNLSIEDTGSSYSNQE---KGED   1562
6   ----------T--SSDI--------TNTKGNFAVK----------------N-------------------NKEIMKTYR   1502
7   --------------------------------------------------------------------------------   1375
8   ----------VSSSTELVPTIVKSVSQQLSTI-PK----------------GSEKADLNCSGNLSTITLSSNDQTMTNKN   1547
9   --------------------------------------------------------------------------------   1427
10  ----------VSLSTEVVRTNVKSVANEESLLTLR----------------NSQKIDPKCSERSLTATLSLDKQTVTEKK   1546
11  ----------S---------------------------------------------------------------------   1387
12  ----------L---------------------------------------------------------------------   1383

    GXXXXXXX---X-XXXXXXIXX--XXXXXXXXXXXXXXKX----------XXXXXXXXXXX-XXXXXXXXXXXXXXX---
1   GKGQVEHVQRQENQESMNKINS--KSFTQDNIAQYKSVKK----------AR-PNNEGENNDYACNVEQASPVRNEVPGD   1610
2   ---------HLP--NNIHEIDI--DKEEN-----TTPKKE-AQMTI--NTQAVNES----N-SATNLEVRLSALPIE---   1514
3   GDKTVNNN------ATTND-------FEY--RTSEDPI----------------------------VSVNIPIFNSR---   1471
4   GF--EERL---PTNQTSQEVVTDLGGERG--SPHAKSVKSNDSIIVQSKRANKNNTNEEPNDIECVEGKSGQA-------   1629
5   GASKSSRS---PHR----EIETHVEDEHV-TVPELNMHK--HSIHIISN--NVTENMEQPRNKTHIEGTTISKEKKGLED   1630
6   GHGDLMTS-----PVRFYEIEK--D--DGIPISVSSPIKR----------QKINFLLPNSN----VIERNL---------   1550
7   ------------------------DGFSLTPVAQAIPVTY----------PVAQTS------------------------   1397
8   GKEQRERTDNLEKQDSINNINK--ESFMQDKITQNEPVKE----------TALQISEDEGKYRTCIVERALPIINAELSD   1615
9   --------------------------------------------------------------------------------   1427
10  GNEQEEKDQSLEKKESMNIIDN--ESPTEDNITQNKTVKK----------TASHNNENEDKNWTYNVEQTLPAMNEGMAD   1614
11  --------------ASQNILKV--QNFKVS---KVSKVKN----------PDVAFI------------------------   1414
12  --------------SSEDAIDV--EDLEPESRRENSNMHS----------PDILTN---------NLET-----------   1417
```

```
   XXXXXXXX-XXXNXXXX-XX-XXXXXXX---XXXX---X--------XXX-----XXXXXXXXXXXXXX-----XXXX---
1  GIQIPSGT-ILLNSSKQ-TEKSKVDDLRSDEDEHGTVAQEKH-QVGAINSRNKNNDRMDSTPIQGTEEE----SREV---  1680
2  -----TGTQMIDNSNKY-------KG-------SEMVTGSDLRAEGKVS--------------------------------  1544
3  -----------------------VSGLD------------------K--------------------------------  1477
4  ------T------------------------------G----------------QSFKANPSKSSAD--SDKPTNLDTK  1654
5  MTENSSI------------------------------G----------------EAKLALPMATIVDDPQSRSTNVDTE  1663
6  --KATQETEISNNGNATDIVPIEVNGDHMDNDGAVSLAGNAT-GGGKNN-----NVVIEAVTPNSESDR----GRHV---  1615
7  -----VEE-ITTNKHKD--------A-----QDPG--------------------------------------------  1413
8  GLQNSSGI-MLVNGLKQ-KEGSKVDTLNHGEDEKVTFAEEDP-QMISVNSSEENAQSIDSVPMQNAVKG----SVEA---  1685
9  --------------------------------------------------------------------------------  1427
10 GIHTSSEA-IFLNNSKQ-TEKSKADALHYNEDEHGMATGENR-QSDEINSVNQNAKKIDSIQIQSVEEE----SRKA---  1684
11 SSTSCPKT-PFTTGSSK-NV-TEINDAV---LE-------------STRSLERGDD----------------------  1451
12 QISSNGKG-SFLNGNGK-AL-LPYDAVR---DA-------------RNRDINENEEH---------------------  1455
```

```
   XXXXXXXXXXXXXXXXXXXXXXXXXXXXXXXXXXXDDFXPXXX------X-XXXXXXLX--XMXXXXSXX---------
1  VMTEEGINVRLEDSGTCELNKNLKGPLKGDKDANINDDFVPVEE------N-VRDEGFLK--SMEHAVSKE---------  1742
2  ---------KTDRSGSII-----------TRNSNVNH--RPGPE--------NSSGLLTLAIYKSTASR---------  1583
3  -----------------------------LNNQKISDEYVISSHDLELARNGLQNQAQLH--DMHEGYKKKSANLLTQ--  1524
4  EMEEDNLKSRNELRTALDEKETKSRTYIEGEEDKGEDDFAPVNT------A--QQDPFLE--AMEKEVSQTEVIL-----  1719
5  ERENSTTSKSSHLIGAL------------EEQQATQDDFVLVEE------P--PKDTLLD--ELDKEISQDMTET-----  1716
6  RISSDSITNKV-------SRNDDFVPVESTRNSSDNDDFVPIEA------A-IEEDNFIE--EMEKEAIRN---------  1670
7  ------------------QKKGANIDKKKEATVESNDDFAPAET------D--QDDEFLL--AMEQKLETTSNKLAPIIG  1465
8  ITTEETLTVHLRDSETTEFNKMMERQQRKGQKINTNDDFAPVDE------N-DRNEGFLE--AMEQVASRE---------  1747
9  --------------------------------------------------------------------------------  1427
10 VTIEESLSERLEGCGTLEPIKNLEGQLKEDKKANINDDFVPVEE------D-GRDEGFLK--AMEQVASKE---------  1746
11 -------F--------APVNVAP------------KDDFLE------------EMEEKIK--IKNQKSSEQ---------  1481
12 ------VFSRI--LGATEFRTFP------------KDEARETKE------VREEKTDNLE--ERPQTVKDD---------  1498
```

```
   -------XXXXXXXEXXXXSXXXXIRIPIFNSXKXXXXXXXXXXXXXXXXX----XX---------------XXXXXX---
1  ------TG-LEEQPEVADISVLPEIRIPIFNSLKMQGSKSQIKEKLKKRLQ-------------------RNELMP---  1792
2  -------------EAPNTQFAPEVHF---GPKKVEPVTSVKMIK---RDQGSMGDIV---------Q----E-------  1623
3  -------HNMRPPFEDVAIPTHKELKIPIFNSLILKSTNQQPMIPNSQRKV-------------------QSEVAI---  1574
4  -------KSSPIAKNKRPTSSMHEINIPIFNTLKLTDPSWKERSRTQQESTTRMGQQKQIDSSQVTST---RS-----D-  1783
5  -------NASTAHKDVVLDSPMPAIRIPIFNTLKIGSGRIEKKKHTKIDKA--LNKRLNLDSSAIILH---QQEDAPCDV  1784
6  -------------LERPAKHHIPSIRIPLFNVSRPSGRENTKQQKDNTHTG----AISTFSDVGVKINEGNFNTIED---  1730
7  QNPPEQNKLQSVSLDDVDAPASSKIRFPIFNSSKFQGRSVKVLNNTTNLTQ----NA-------------K---------  1519
8  ------TSLLKEQTELADTSVLPEIRIPIFNSLKIQESNGQIKQKLKMRLQ-------------------RNESMP---  1798
9  --------------------------------------------------------------------------------  1427
10 ------TSLLKEQPEVADISVLPEIRIPIFNSLKIQESKSQIKEKLKKRLQ-------------------RNELMP---  1797
11 --------IEGIEANHNDVSQMSQIRIPIFNSSKLQNKLDLEGPRCDKGE--------------------------  1523
12 --------DAMASVENKDESQVLGIRIPIFNSSKLQNKPKSKDSVHENDLS----ER-------------KRKRVDG---  1550
```

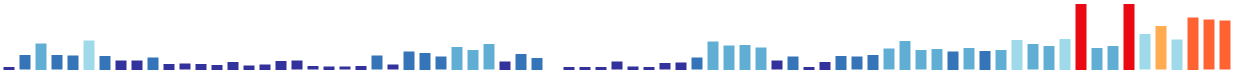

```
    ----XDXXPXXXX----X-XXX----XXXXXXXXXX-------XXXEXXLXX-XXXXXXXXXXXXXXXDXXSXEXTPS

1   ----PDSPPRMTENTNINAQNGLDTVPKTIGGKE-KH---------HEIQLGQAHTEADGEPLLGGDGNEDATSREATPS   1858
2   ---EKDVNTAFEP---FQQDAAIAKV-----------ALKDDEMSCEIQ-----------VSENENCADDTSSKEGTPS   1674
3   -----------------R-KR-----PLGNPSTQGQY--------------------TDIENERVRDYDDSQSLEDSAD   1610
4   ---KQDVIP-----------------------------------SHLEVDLHS--SFPYGTPGNGIDDDMDINSRETTPS   1823
5   PIITKDHLPEEIDTVT-KQAPAFGNIPVNNMAEE----------VPLENDLQS--SFVIDG-----KEDGDGTSRDTTPS   1846
6   ----KKMNPSFNESTEVL--AG----KTNSKGMEARKDEFKDDIEDDEIE-----SSSSEDRMQDDEEYRDEASSASTQS   1795
7   ----------------------------NADEKEPDN-----------------NARSNGQTLEESRCSEDAVSRDATPS   1554
8   ----PDSPPRVASNANTNNVLN----DSTSVGKEGKY---------DEIQLAQSHIEKDGDPLLVGEGNEDATSREATPS   1861
9   --------------------------------------------------------------------------------   1427
10  ----PDSPPRLAANTNIIGRNGLGTAPKAIGGKEEKH---------GEIQLAQAHTEADADPSLVGDGNEDATSREATPS   1864
11  ------ESPREPR-------------D-SSVGKACDK----DAKNSQAIDLES-SQEDNESNILDNNSKEDY-SKEPTPI   1577
12  ----NDEEGNDYD-------------E-HEVETAREV----DPISTQPEDLAS-S-PDNESTFLQDSRRSDWASQNSAPS   1606
```

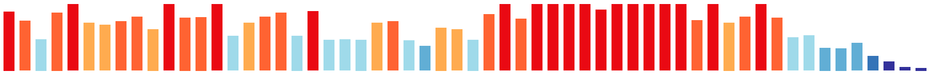

```
    LRXHFPSKKXRKLVXRLRXFXXXDLXXXSXEEXRNLRIELLDFMMKLEYXXXXXXX----

1   LKVHFFSKKSRRLVARLRGFTPGDLNGISVEERRNLRIELLDFMMRLEYYSNRDNDMN--   1916
2   LHIHFPSKKARKIVNRLRDFKPDQLAALSAAERRNLRVELLDFMMKMEYFEAEE------   1728
3   IRLHFPNKRARRIVSKLRGFSEYDMGQLSAEEKRNLRIELLDFLMKIEYHTITD------   1664
4   LRMFFPSRKTRRLVSRLRNFDPSELSTLPTEERRNLRIELLDFLMKLEYYTGDDADKREI   1883
5   LRMHFPCRKSRKLVNRLRTFNPDELASLPVEERRNLRIELLDFMMKLEYYTGDKNN----   1902
6   LRDHFPSQNARKLVRYINELGHEELLQFLPEEKRNLRIELLDFLMKLEHYSDK-------   1848
7   LRMHFPSKKARKLVNRVRAFTAEDLSHLPPEEKRNLRIELLDFLMSLEHEHGFYQ-----   1609
8   LKIHFSSKKSRKLVSRLRGFTPGDLNGISVEERRNLRIELLDFMMRLEYYSNRDDDMV--   1919
9   ------------------------------------------------------------   1427
10  LKAHFSSKKSRRLVGRLRGFTPGDLNGISVEERRNLRIELLDFMMRLEYYSNRDNDMG--   1922
11  LRLHFPSKKARKLVSRLRGFTVEDVSDISPEEKRNLRIELLDFMMKLEHESLNGEF----   1633
12  LRLHFPTRKSRKLVSKLRGFSLDDISRISLEEKRNLRIELLDFMMKLEHESLAEEL----   1662
```

**Consensus Threshold:** >50%

**Compare to:** Saccharomyces_cerevisiae

Amino acids that match the reference are marked with orange highlighting.

**Created:** Aug 10, 2020

**Last Modified:** Aug 10, 2020
